# High-contrast en-bloc staining of mouse whole-brain samples for EM-based connectomics

**DOI:** 10.1101/2022.03.30.486341

**Authors:** Kun Song, Zhihui Feng, Moritz Helmstaedter

## Abstract

Connectomes of human cortical gray matter require high-contrast homogeneously stained samples sized at least 2-3 mm on a side, and a whole-mouse brain connectome requires samples sized at least 5-10 mm on a side. Here we report en-bloc staining and embedding protocols for these and other applications, removing a key obstacle for connectomic analyses at mammalian whole-brain level.

## MAIN TEXT

The dense and homogeneous deposition of heavy metals into brain tissue that leads to high membrane contrast for electron-based imaging is a prerequisite for synaptic resolution connectomics. Ever since the development of the “reduced-osmium” protocols (Bruijn, 1973; Karnovsky, 1971; Willingham & Rutherford, 1984), *en-bloc* staining of tissue samples up to a thickness of about 100-200 μm was possible. Beyond such sample sizes, however, substantial staining gradients occurred, which limited connectomic analyses to smaller samples (Briggman, Helmstaedter, & Denk, 2011; Holcomb et al., 2013). With the development of a modified staining protocol (Hua, Laserstein, & Helmstaedter, 2015), samples up to about 1 mm in size could be homogeneously stained – an important step to allow millimeter-size connectomic data acquisition (Fig. 1a). This protocol has been widely applied since (Kenneth J. Hayworth et al., 2020; Kuan et al., 2020; Alessandro Motta et al., 2019; Yin et al., 2019).

**Figure 1.**
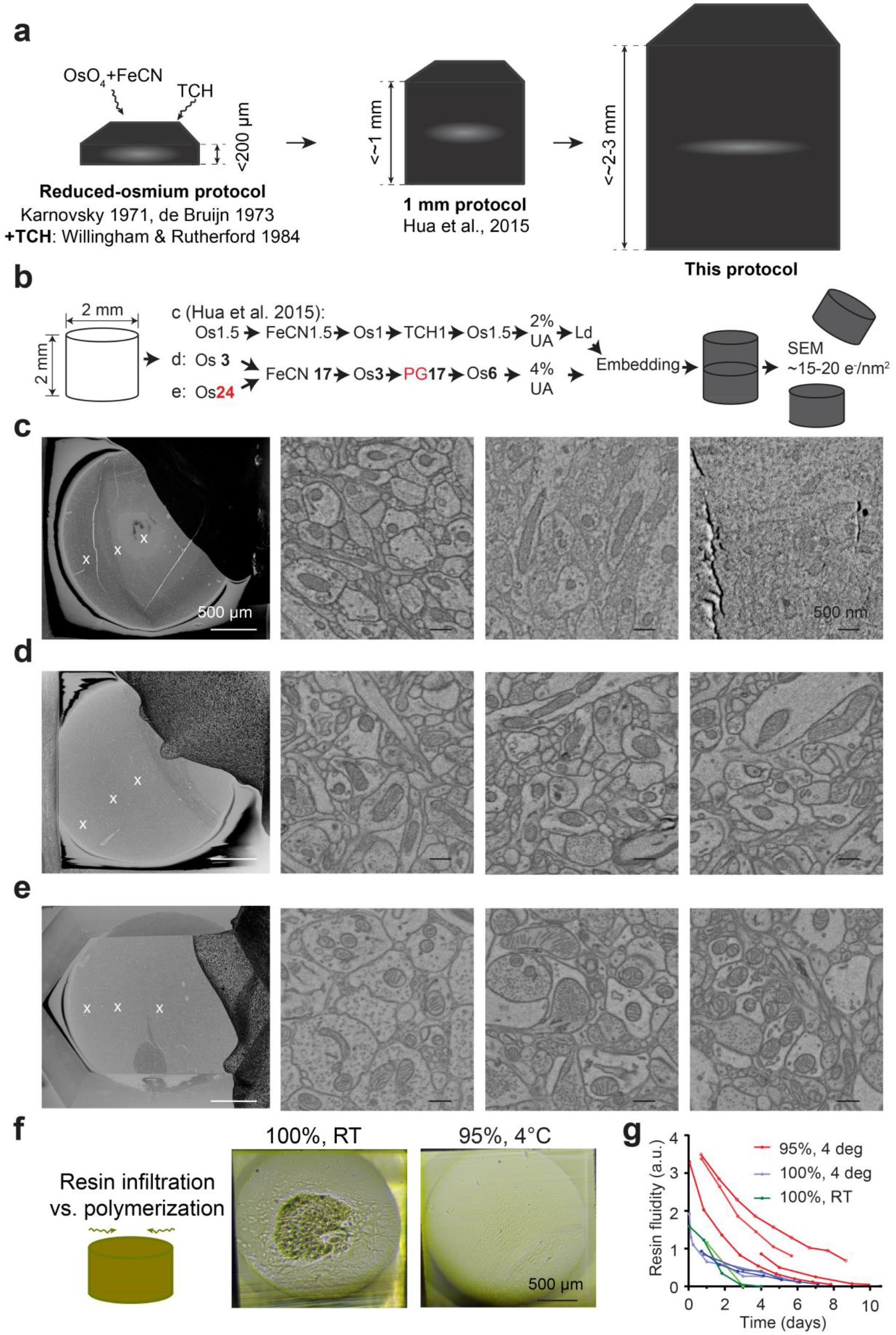
High-contrast staining protocol for 2-3 mm sized samples. **a,b** Staining gradients were the key limitation to sample size for decades. The initial reduced osmium-TCH protocols(Bruijn, 1973; Karnovsky, 1971) yielded high-contrast homogeneous staining for samples up to about 200-250 μm thickness. Heavy metal atomic nuclei were provided mainly by Osmium and Uranium compounds, with enhancement by thiocarbohydrazide(Seligman, Wasserkrug, & Hanker, 1966) and ferrocyanide(Bruijn, 1973; Karnovsky, 1971; Willingham & Rutherford, 1984). A modified protocol(Hua et al., 2015) allowed the staining of up to 1 mm sized samples. The protocol reported here expands this to 2-3 mm sized samples, large enough to cover the depth of the human cortex. **c** Staining of 2mm sized sample using the(Hua et al., 2015) protocol yields staining gradient beyond 1mm with low-contrast core. From left to right: overview image showing three layers of gradient; high-res EM image of outer position in sample, note the contrast was comparable to 1 mm samples; middle position shows sharply decreased contrast; in center, no meaningful images can be acquired due to low signal. **d,e** This protocol removed these gradients. See Table 1 for protocol steps and their relevance; note even better contrast after prolonged initial OsO_4_ incubation length (e). **f,g** Resin infiltration speed in dependence of temperature. Resin infiltration of larger samples requires separation of diffusion from polymerization to avoid softer core of sample (f). With appropriate infiltration protocol (Suppl. Table 1), a softened core can be avoided.

However, with the ambition to obtain connectomes from even larger samples, in particular samples that encompass the gray matter depth of human cortex (2-3 mm in size), and samples corresponding to entire brains of small mammals like mouse (Abbott et al., 2020) or even humans (Motta, Schurr, Staffler, & Helmstaedter, 2019), the need for improved protocols became obvious. In spite of the promising initial attempts for mouse whole-brain staining (Mikula & Denk, 2015) there is so far no reliable protocol for the en-bloc staining of 2-3-mm-to-centimeter scale samples with high staining contrast. This is particularly challenging since EM imaging and reconstruction have progressed to a stage at which large-sample analyses appear to become possible(Kenneth J. Hayworth et al., 2020; Januszewski et al., 2018; Alessandro Motta et al., 2019).

Here, we report such protocols for 2-3 mm-sized samples (Fig.1), mouse hemispheres (Fig.2) and mouse whole brains (Fig.3). Their development required the concomitant solution of the following problems: recurring staining inhomogeneity, sample instability that leads to breakages (especially in hemispheres and whole-brain samples), and homogeneous resin infiltration, as described in the following.

**Figure 2.**
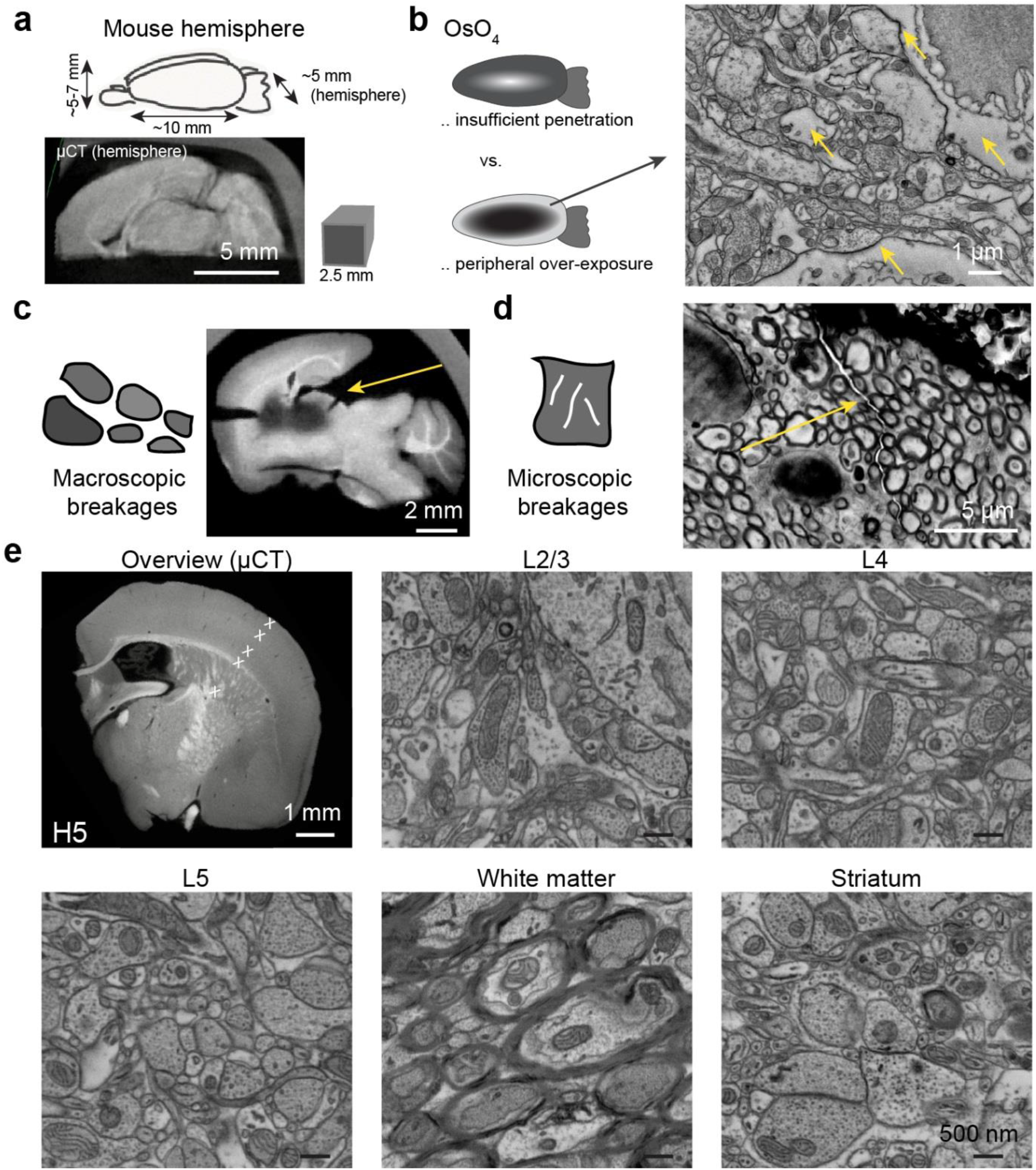
High-contrast staining protocol for mouse brain hemispheres. **a** Size of mouse brain hemisphere compared to the 2-3mm block size reported in Fig. 1, and μCT image of mouse hemisphere as used for protocol development. **b-d** Key challenges of hemisphere and whole-mouse brain staining were sufficient infiltration of OsO4 without over-exposure of OsO4 in the periphery, that contains the very relevant gray matter of the cerebral cortex (b); breakages of the sample at macroscopic (c) or microscopic (d) scale. Note only the macrosopic breakages could be clearly identified in μCT, but the microscopic were as destructive to the goal of whole-brain connectomic analysis. **e** Homogeneously stained mouse brain hemisphere using protocol developed here (see Table 1), note high contrast from cortical to subcortical positions (approximate locations of high-res EM images indicated in μCT overview).

**Figure 3.**
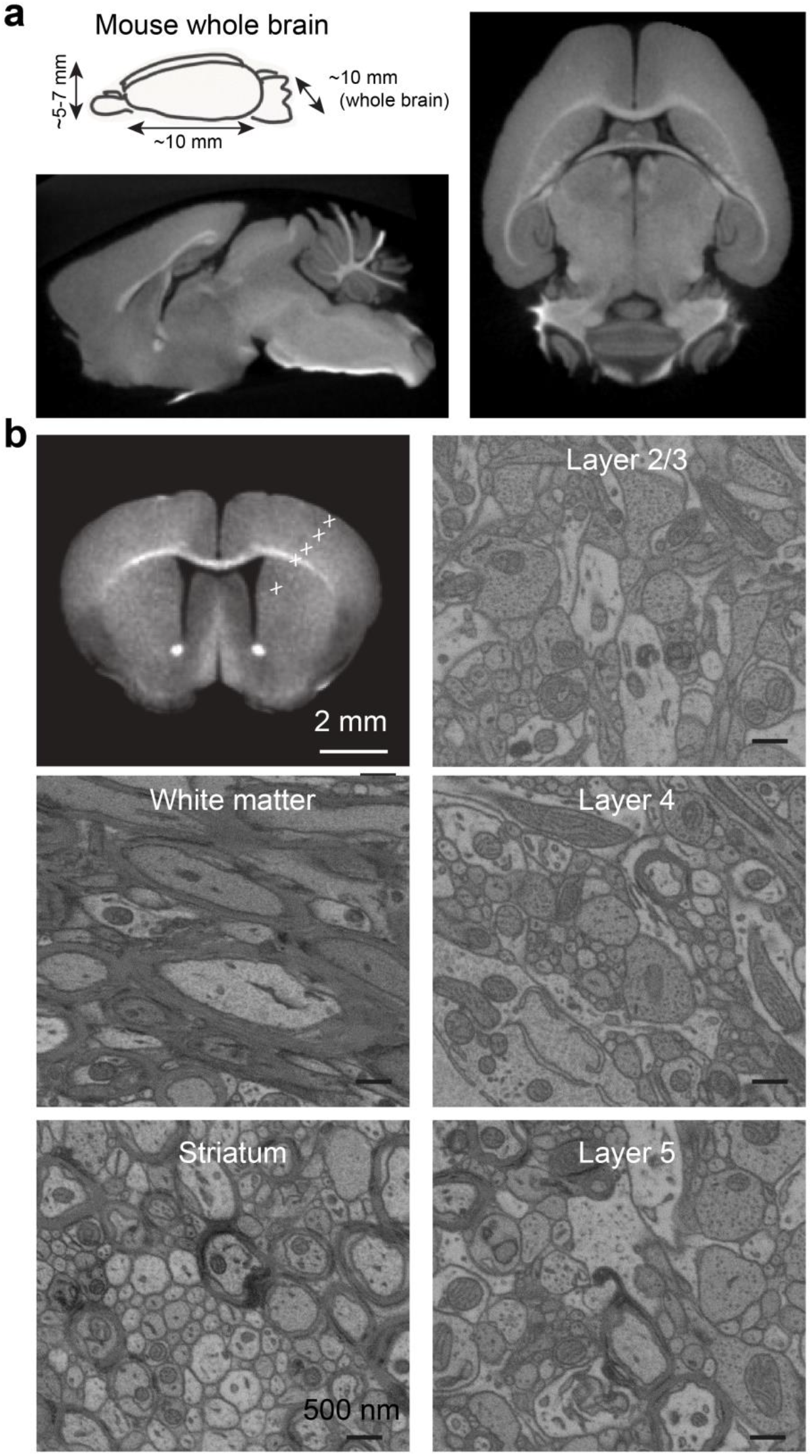
High-contrast staining protocol for whole mouse brains. **a** Size of whole mouse brain, and μCT cross-sections of fully and homogeneously stained whole mouse brain (sagittal plane, bottom; transversal plane, right) using the protocol developed here (Table 1). **b** High-resolution EM images acquired at various positions along the mouse brain from superficial cortex to subcortical tissue, approximate positions indicated in μCT overview (top left).

For protocol development, we initially used X-ray microtomography (μCT) imaging to assess staining gradients (Mikula & Denk, 2015; Ströh, Hammerschmith, Tank, Seung, & Adrian, 2021). Additionally, we applied low-vacuum SEM (Kazmierski & Millington, 1979; Sommons & Marquis, 1997) to check ultrastructural contrast after intermediate steps of the staining experiments, at which stages the sample would be charging in high-vacuum SEM. This was of particular importance as samples that had homogeneous appearance in μCT could reveal insufficient membrane contrast or damaged ultrastructure when analyzed in EM (Supplementary Figs.5, 7).

First we applied the available 1-mm protocol (Hua et al., 2015) to 2 mm-sized samples (Fig.1b), which however yielded strong staining gradients (Fig.1c, Suppl.Fig.1a,b) and incomplete resin infiltration (Fig.1f). We then used μCT to investigate at which steps in the protocol the gradients occurred and found a two-layered gradient after OsO_4_ - FeCN incubation, and an additional third gradient layer after thiocarbohydrazide (TCH)-OsO_4_ incubation (Supplementary Fig.1c). The obvious measure when observing such gradients was to extend the duration of the respective incubation steps. When extending the FeCN step from 1.5h to 12-17h, the gradient was in fact removed (Supplementary Fig1d). To our surprise, however, when omitting the FeCN step entirely and only incubating in OsO_4_ for 3h, membrane contrast under SEM was not reduced (and there was no gradient) (Supplementary Fig. 2a). Only when we also extended the OsO_4_ incubation to 24h, the FeCN step provided enhanced membrane contrast (Supplementary Fig. 2b; for a possible explanation for this phenomenon by means of Os(vi)-CN^−^ coordination chemistry, see Suppl. Material and Supplementary Fig.6d,e).

While the FeCN-induced gradient could thus be removed, the TCH-induced gradient could not be similarly resolved; rather, prolonged TCH incubation yielded broken samples, most likely because of gaseous products generated by the reaction of OsO_4_ and TCH as reported previously(Mikula & Denk, 2015) (Supplementary Fig.1e).

We therefore adopted the exchange of TCH by pyrogallol (Pg) as proposed in (Mikula & Denk, 2015). When replacing TCH by Pg in the 1 mm protocol (Hua et al., 2015), we found sufficient sample conductivity and contrast (Supplementary Fig.1f,g,h). We confirmed that pyrogallol incubation should be performed in H_2_O instead of cacodylate buffer (CaC), which would otherwise cause decreased membrane contrast(Mikula & Denk, 2015) and resulted in a staining gradient (Supplementary Fig.1i). In addition, we found it necessary to pre-incubate in H_2_O to wash out CaC before the pyrogallol step, since interaction of CaC with pyrogallol could possibly cause gradients or breakages (Suppl. Fig. 3g). By extending the incubation times for pyrogallol and the following OsO_4_ step, the second gradient vanished (Table 1, Supplementary Fig.1j).

**Table 1.**
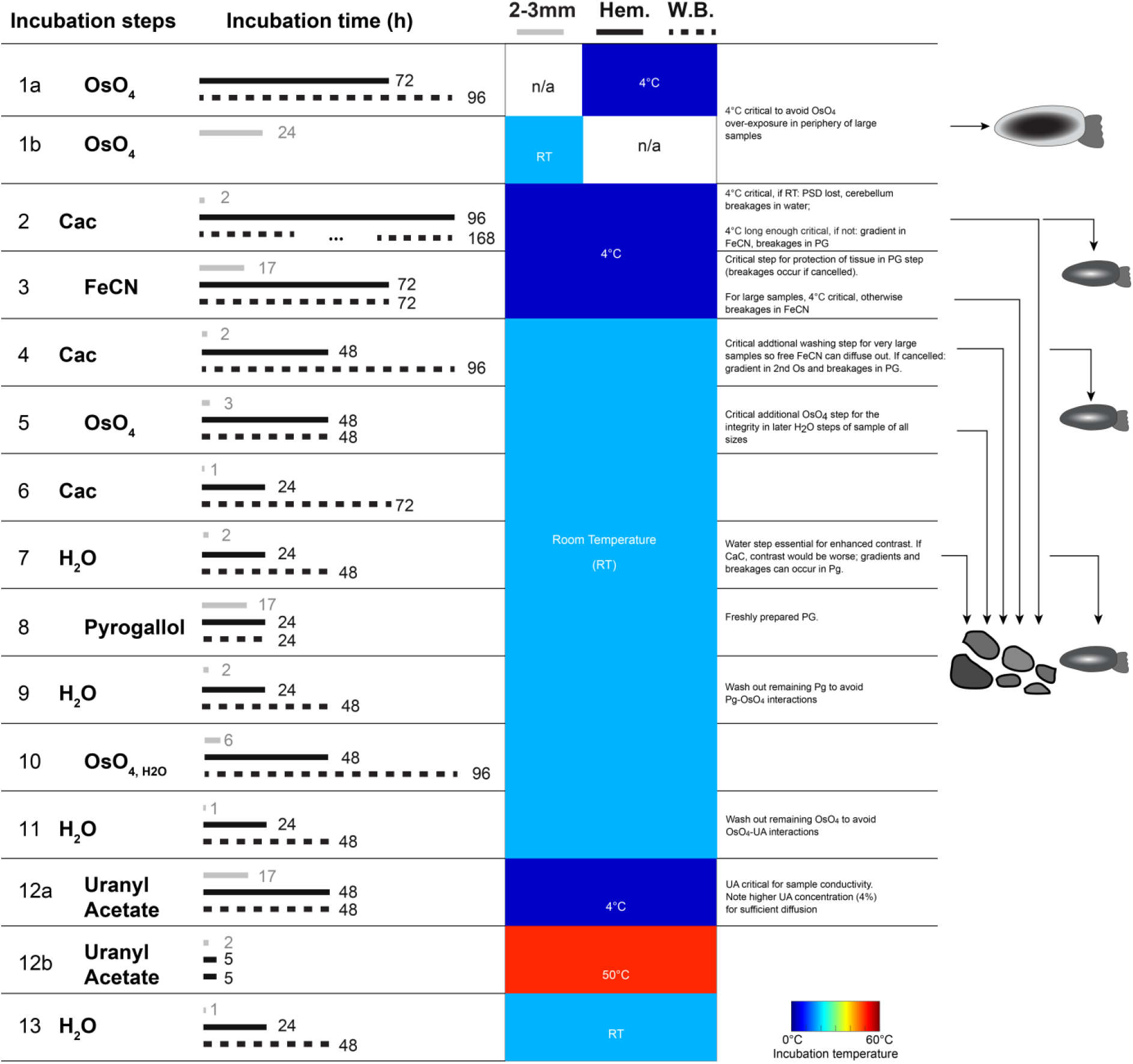
Overview of 2-3mm, hemisphere and whole-brain protocol steps with incubation times and temperature steps visualized. Illustration of the relevance of various staining and washing steps for the key challenges of whole-brain staining: avoidance of gradients, peripheral tissue destruction, and macro- or microbreakages.

For the final steps in our protocol, uranyl acetate (UA) and lead aspartate (Ld), we noticed under μCT a minor gradient caused by UA (Supplementary Fig.1k). This gradient was removed by increasing the UA concentration from 2% to 4% (Supplementary Fig.1k). We omitted the final lead aspartate step as the low concentration of lead yielded very bad diffusibility (20 mM/L lead nitrate dissolved with 2:3 mole ratio in 30 mM/L saturated aspartate buffer (Walton, 1979)) (Supplementary Fig.1k). Also under omission of the lead aspartate step, sample conductivity and staining contrast were sufficient (Fig.1d,e). With this, we had obtained a high-contrast homogeneous staining protocol for 2-3 mm sized samples (Fig. 1e, Table 1).

Next, we needed to ensure homogeneous resin infiltration. We focused on Spurr’s resin for samples to be cut and imaged using SBEM(Denk & Horstmann, 2004) and Epon 812 substitute (in the following referred to as Epon) for samples to be cut using ATUM(K. J. Hayworth, Kasthuri, Schalek, & Lichtman, 2006). We considered the fact that the epoxy resin blender would undergo polymerization during infiltration. This would increase the viscosity of the blender; and once the polymerization process has crossed the gel point, no more diffusion would be possible(Vidil et al., 2016). Thus the practical strategy for improving resin infiltration was to slow down polymerization reactions and keep viscosity low. To do so, we kept all resin infiltration steps at 4°C(Vidil et al., 2016) which in fact significantly slowed down the polymerization process (Fig.1g). Furthermore, before incubating the samples in pure resin, we added a step of 95% resin in 5% acetone; this small amount of acetone was shown to significantly decrease the resin viscosity(Loos, Coelho, Pezzin, & Amico, 2008). We finally extended the resin (Epon) incubation time from ~1 day to 4-5 days (Fig.1g; Suppl. Table 1, which also contains data for Spurr’s resin). By these modifications, we successfully infiltrated both Spurr’s and Epon epoxy resin into the center, yielding homogeneously embedded 2 mm-sized samples (Fig.1f).

Next, we wanted to apply the obtained protocol to the staining of mouse hemispheres and whole-brain samples that are about 25-60 fold larger in volume (about 5×10×(5 or 10) mm^3^, Fig. 2a, 3a). For protocol development, we used mouse brain hemispheres as a surrogate goal (Fig. 2), since the mouse whole brain is a symmetric duplication of these, with nominally similar access geometry to the hemisphere, and later tested our results in an exemplary whole-mouse brain staining (Fig. 3).

Since we had noticed a strong contrast enhancement from prolonged OsO_4_ incubation already for 2-3 mm sized samples (see above, Fig.1d,e, Supplementary Fig.2a,b), for mouse hemispheres, we first extended the initial OsO_4_ incubation to 3 and 6 days. In fact, OsO_4_ incubation for 3 or 6 days resulted in increased membrane contrast, also in the sample center. But we encountered a serious obstacle: when incubating for sufficient duration for OsO_4_ to reach the center of the hemisphere sample, the outer parts of the sample were showing strong signs of ultrastructural disintegration (Fig.2b, of note, these were not apparent in μCT, only under EM imaging). The cytosol of most neurites and cell bodies had an extracted appearance, possibly indicating the removal of intracellular proteins (Fig. 2b, note that Hayat 1981 had already discussed this in the context of over-fixation of tissue by OsO_4_). Since the outer part of the sample corresponds to the cortical gray matter, a key target of whole-brain connectomic analysis, a whole-brain staining with good stain penetration but insufficient ultrastructural quality in the cortical periphery would be useless.

In order to overcome this obstacle, we took the notion that the ultrastructural alterations were likely a result of protein over-oxidization by prolonged OsO_4_ incubation (see Supplementary Material), which could be slowed down by lower incubation temperature (4°C) (Hayat, 1981). In fact, no obvious OsO_4_-based cytosolic alterations were found for samples incubated at 4°C, even when incubating in OsO_4_ for 7 days (Supplementary Fig.2e,f). With sufficiently long OsO_4_ incubation at 4°C (step length was diffusion time plus about one additional day incubation for hemispheres), a step to RT was no more necessary, and the subsequent FeCN still yielded enhanced membrane contrast. (Suppl.Fig.2g,h,i,j; see Supplementary Material for possible chemical explanation, Supplementary Fig.6d,e). The FeCN incubation itself also induced substantial tissue damage at RT (Supplementary Fig2.g). We therefore incubated FeCN at 4°C, as well, which resolved this problem (Supplementary Fig2.j).

For the FeCN step, we noticed an additional issue: especially for larger samples such as the hemisphere, we observed an inverse gradient with lower μCT-imaged intensity in the periphery (Supplementary Fig.3b), and a very slow diffusion of FeCN (Supplementary Fig.3d). In our current understanding, this could be the result of the direct redox reaction between OsO_4_ and FeCN (see Suppl. Material). When however adding washing steps with CaC between OsO_4_ and FeCN incubations to avoid their direct interaction, the ensuing gradient could be removed, staining intensity in the periphery remained high (Supplementary Fig.3c), and the FeCN diffusion was much faster (Supplementary Fig.3e,f). We noticed that these washing steps should be at 4°C and of sufficient duration (Supplementary Fig.3h); if they were not sufficiently long, there remained a staining gradient that would be amplified in the pyrogallol step later and may contribute to breakages (Table 1, Supplementary Fig.3g). If they were at RT, while the incubation time could be shorter (Supplementary Fig.3h), the samples would be less stable in water with tissue damage in cerebellum, and we also noticed a reduction in PSD staining (Supplemenatry Figs.4, 7f-g)

After having addressed the OsO_4_ penetration issues, we were left with one major remaining obstacle: larger samples were consistently broken during the staining process, either in their entirety into several smaller pieces, or with micro-breakages that would hamper dense circuit reconstruction (Fig. 2c,d). These breakages would usually occur in relation to the pyrogallol incubation step (Table 1). We therefore had to assess the effect of all earlier protocol steps on the stability of the sample during pyrogallol incubation. This was of particular importance since the pyrogallol incubation (including the washing steps before and after) had to be performed in H_2_O, as determined for the 2 mm samples; and this H_2_O incubation could induce substantial osmotic forces in the sample. We found that inserting an additional extended OsO_4_ incubation step after FeCN would stabilize samples such that they could be incubated in H_2_O for up to 100 h without the occurrence of major breakages (Supplementary Fig. 3i, a possible explanation is the membrane perforation induced by OsO_4_, which increases tolerance to osmotic stress, Hayat 1981). While this additional OsO_4_ step may also serve to enhance background staining and thus sample conductivity for SEM, in our view the enhanced stability of the large samples is the most critical aspect.

Finally, we applied the hemisphere protocol to n=3 hemispheres (Fig. 2, Supplementary Fig. 4), which all remained intact and provided homogeneous high-contrast staining throughout the sample. In addition, we stained an entire mouse brain with modest additional extensions of incubations times (Table 1, Fig. 3), yielding a whole-mouse brain staining protocol for large-scale connectomics (Abbott et al., 2020). We observed two types of artifacts that remained in large-sample staining (Suppl. Fig. 7): the detachment of larger blood vessel walls from the surrounding neuropil, and rare remaining microbreakages in the subcortical regions with high rate of myelinated fibers. Also, special care had to be taken to preserve the integrity of the cerebellum (Suppl. Fig. 7f,g). Neither of these remaining issues are expected to substantially affect connectomic reconstruction.

We envision that the protocols reported here will serve for large-scale connectomic projects in mouse and other species. In particular, the volume of a mouse brain also approaches relevant cortical and subcortical volumes in higher mammals as non-human primates and humans. The ultimate goal of obtaining large connectomes from fractions or the entirety of human cortex(A. Motta et al., 2019) will also profit from the advances described here, both for fundamental research and clinical applications.

## ACKNOWLEDGEMENTS

We thank Yunfeng Hua for sharing very helpful insights into staining protocols, for discussions and advice, Erin Cocks for contributions to initial experiments and discussions, Meike Sievers for initial contributions to resin protocols and comments on the manuscript, and Niansheng Ju for contributions to control experiments. We are also grateful for brief discussions of chemical topics with Daniel M. E. van Niekerk, Chao Sun, Shi Chen, Thomas S. Hofer; to Martin Kind, Marcus Koch and Thaleia Vavaleskou for support and advice on EDS experiments; to Markus Braun, Chokri Boumrifak and Yagmur Aydogan for support with UV-vis spectrum and Raman spectrum measurements. We thank Smaro Soworka, Selina Horn, Lev Dadashev and Iris Wolf for excellent technical support.

## METHODS

### Animals Experiments

All experimental procedures were approved by the local animal care and use committee and were in accordance with the laws of animal experimentation issued by the German federal government (Regierungspräsidium Darmstadt, Germany, permits V54 - 19 c 20/15 - F126/1028 and F126/1002).

Adult C57BL6/J mice (male/female, P30-P90) were treated with analgesics (0.1 mg/kg Buprenorphin (CP-Pharma) and 100 mg/kg Metamizol (WDT)) 0.5 h before isoflurane anesthesia (Harvard Apparatus, 5% in O_2_ for initialization, 2-3% for maintenance, O_2_ flow rate 1L/min). Following anesthesia, the animals were transcardially perfused (Harvard Apparatus, flow rate 10 ml/min) using 15 ml sodium cacodylate buffer (0.15 M, pH 7.4, Sigma-Aldrich) followed by 30 ml fixative containing 2.5% paraformaldehyde (Sigma-Aldrich), 1.25% glutaraldehyde (Serva), and 2 mM calcium chloride (Sigma-Aldrich) in 0.08 M sodium cacodylate buffer (osmolarity about 700-800 mmol/Kg, pH 7.4). The duration between start of perfusion and incision of diaphragm was less than 30 seconds. After perfusion animals were decapitated, the skull was opened with care to avoid mechanical damage to the brain, and the brain was post-fixed in situ for 12 to 96 h at 4 °C before extraction from the skull.

For 2 mm samples, the brain was cut into 2 mm thick coronal sections in 0.15 M cacodylate buffer using a vibratome (Leica VT1200). Then, a 2 mm diameter biopsy punch (KAI Medical, Honolulu, USA) was used to extract samples from dorsal cortical, ventral cortical and subcortical regions (Supplementary Fig.1j). The samples were then stored in 0.15 M cacodylate buffer at 4 °C for 8-24 h before staining.

For hemisphere samples, brains were cut with a razor blade (Wilkinson) along the midline. The hemispheres were stored in 0.15 M cacodylate buffer at 4 °C for 24 h before staining.

### Staining Experiments

All staining and resin infiltration steps for 2-3 mm samples were carried out in 2 ml Eppendorf tubes and for hemisphere or whole brain samples in 50 ml glass tubes at room temperature (~20-23°C) unless specified otherwise. For steps involving photosensitive chemicals (FeCN, TCH, Pg, UA or Ld), the tubes were covered with aluminum foil.

All chemicals used in the staining pipeline are listed in Supplementary Table 2; all experiments reported in main and supplementary figures are detailed in Supplementary Table 3.

For simplicity, the following terms were used to refer to the recurring staining steps in different experiments:

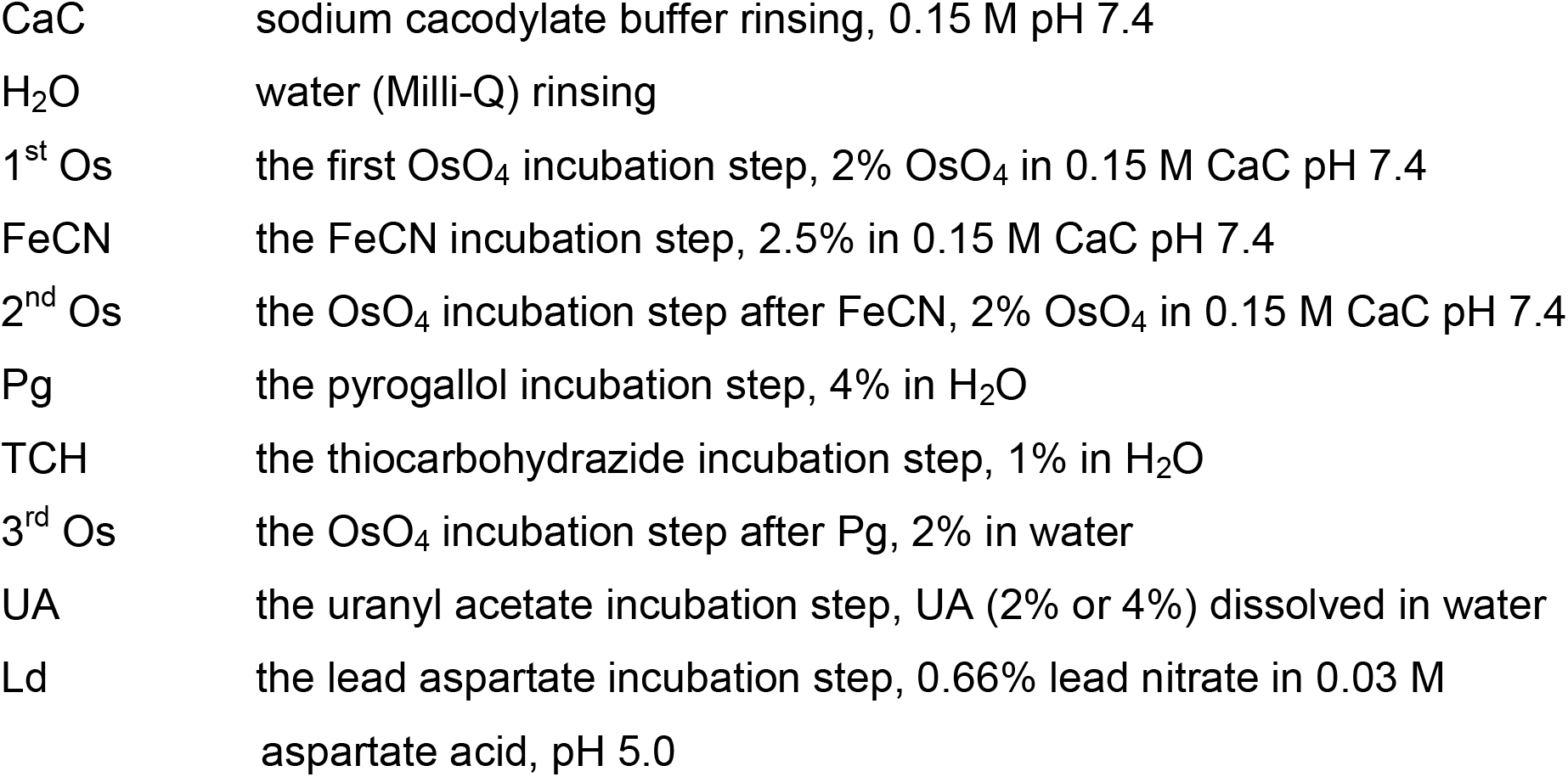

### Staining of 2 mm samples with 1 mm protocol

The 1 mm staining protocol (ref. (Hua et al., 2015) with addition of 2^nd^ Os step) was applied to 2 mm samples directly. The samples were stained with the following steps: 1^st^ Os 1.5 h → FeCN 1.5 h →2^nd^ Os 1 h→CaC 0.5 h →H_2_O 0.5 h→TCH 1.5 h→H_2_O 0.5 h (two times)→3^rd^ Os 1.5 h→H_2_O 0.5 h(two times)→2% UA (17 h at 4°C, 50 h at 50°C)→H_2_O 0.5 h (two times)→Ld 2 h at 50°C→ H_2_O 0.5 h(two times). Afterwards, the samples were incubated in graded ethanol series from 50% (4°C), 75% (4°C) to 100% (each step 45 min). Then the samples were incubated in pure acetone for three times, each time 45 min. Afterwards, they were incubated in 50% Spurr’s resin (Sigma-Aldrich, in the ratio of 0.95 g ERL 4221, 5.9 g DER 736, 0.1 g NSA, 113 ul DMAE) in acetone for 6 h. Then the samples were left overnight with the cap of the Eppendorf tubes open to allow evaporation of acetone. Afterwards, the samples were transferred into pure Spurr’s resin for 6 h before embedding and curing at 70°C for 1-3 days. After resin curing, the samples were first imaged in μCT for staining homogeneity. Then they were trimmed to expose the sample center, the sample surface was smoothed and imaged in high vacuum SEM and EDS.

### Step-by-step μCT diagnosis of the main staining steps of the 1 mm protocol on 2 mm samples

The 1 mm staining protocol (ref. (Hua et al., 2015) with addition of 2^nd^ Os step) was applied to a batch of 2 mm samples as described above: 1^st^ Os 1.5 h → FeCN 1.5 h →2^nd^ Os 1 h→CaC 0.5 h →H_2_O 0.5 h→TCH 1.5 h→H_2_O 0.5 h (two times)→3^rd^ Os 1.5 h→H_2_O 0.5 h (two times)→2% UA (17 h at 4°C, 50 h at 50°C)→H_2_O 0.5 h (two times)→Ld 2 h at 50°C→ H_2_O 0.5 h(two times). After each main step (1^st^ Os, FeCN, 2^nd^ Os, 3^rd^ Os, UA), 2 samples were taken out from the staining pipeline, and rinsed with either CaC or H_2_O depending on the solvent condition of the corresponding staining step (rinsing solution changed every 8/17 h). When the rinsing was done for all samples from different conditions, samples were embedded in 2% agarose in water in the same Eppendorf tube at different tube depth, and stored in 4 °C for the agarose to cure. Then the tube containing all the samples were imaged in μCT to investigate staining gradients.

### Extending FeCN incubation time for 2 mm samples

2 mm samples were stained as follows: 1^st^ Os 3 h → FeCN. During FeCN incubation, 2 samples were taken out at each of the following time points:1.5 h, 3 h, 7 h and 17 h. Samples were then rinsed in CaC for 1 h and embedded in 2% agarose as described above. After agarose curing, they were imaged in μCT to investigate staining gradients possibly related to FeCN incubation.

### Interaction between OsO_4_ incubation duration and FeCN incubation

2 mm samples were stained with the following two conditions: (1) 1^st^ Os **3h** → FeCN 17 h; or (2) 1^st^ Os **24 h** → FeCN 17 h. Then they were rinsed sequentially by CaC (for 0.5 h) and H_2_O (two times, each time 0.5 h); they were then dehydrated and embedded according to the 2 mm Spurr’s resin protocol. After resin embedding, they were trimmed to expose the center, their surface was smoothed and they were imaged in low-vacuum SEM to investigate membrane contrast.

### Extending TCH incubation for 2 mm samples

Three groups of 2 mm samples were stained, each batch was incubated with a different TCH incubation length, using the following protocol: 1^st^ Os 3 h → FeCN 17 h → 2^nd^ Os 3 h → CaC 0.5 h → H_2_O 0.5 h (two times) → TCH for 1.5 h, or 3 h, or 5 h → H_2_O 0.5 h (two times) → 3^rd^ Os 3 h. Afterwards, they were rinsed and embedded in 2% agarose for imaging in μCT.

### Replacing TCH by pyrogallol

Three groups of 1 mm samples were stained, each group differed in the TCH-related step, using the following protocol: 1^st^ Os 1.5 h → FeCN 1.5 h →2^nd^ Os 1 h→CaC 0.5 h →H_2_O 0.5 h→ TCH 1.5 h, or Pg 1.5 h, or H_2_O 1.5 h →H_2_O 0.5 h (two times)→3^rd^ Os 1.5 h→H_2_O 0.5 h (two times)→2% UA (17 h at 4°C, 50 h at 50°C)→H_2_O 0.5 h (two times)→Ld 2 h at 50°C→ H_2_O 0.5 h(two times). Afterwards, they were dehydrated and resin embedded according to (Hua et al., 2015). After resin curing, the samples were trimmed to expose the center, their surface was smoothed and they were imaged in high vacuum SEM and EDS. For EDS measurement, we selected point measurement in the neuropile of the sample center.

### Comparison of pyrogallol incubation in H_2_O vs. CaC

Two groups of 2 mm samples (the groups differ in the Pg incubation and the water steps around Pg) were stained with the following sequential incubation steps: 1^st^ Os 24 h → FeCN 17 h → 2^nd^ Os 3 h → CaC 0.5 h → H_2_O 0.5 h (twice) → Pg in water 17 h → H_2_O 0.5 h (twice) or CaC 0.5 h (twice) → Pg in CaC 17 h → CaC 0.5 h (twice)→ 3^rd^ Os 6 h → H_2_O 0.5 h (twice) → 4% UA (17 h at 4°C, 2 h at 50°C) → H_2_O 0.5 h (twice). Samples were then dehydrated and embedded in Spurr’s resin as described above for 2 mm samples. After resin curing, they were trimmed to expose the center, their surface was smoothed and they were imaged in high-vacuum SEM.

### Long-duration OsO_4_ incubation on 2 mm samples

2 mm samples were stained with 2% OsO_4_ in CaC for either 3 days or 6 days. After rinsing by 0.15 M CaC for 0.5 h and two times water (each time 0.5 h), they were dehydrated and embedded according to the 2 mm Spurr’s resin protocol. Then they were trimmed to expose the center, their surface was smoothed and they were imaged in low vacuum SEM for determining ultrastructural preservation.

### Extended Pg and 3^rd^ Os incubtation for 2 mm samples

Three groups of 2 mm samples (groups differ in the incubation duration of Pg and 3^rd^ Os steps) were stained using the following protocol: 1^st^ Os 3 h → FeCN 17 h → 2^nd^ Os 3 h → CaC 0.5 h → H_2_O 0.5 h (twice) → Pg in water (for 6 or 17 h) → H_2_O 0.5 h (twice) → 3^rd^ Os (for 3 or 6 h) → H_2_O 0.5 h (twice) → 2% UA (17 h at 4°C, 2 h at 50°C) → H_2_O 0.5 h (twice). The combination of Pg and 3^rd^ Os incubation time were as follows: (1) Pg 6 h - 3^rd^ Os 3 h; (2) Pg 17 h-3^rd^ Os 3 h; (3) Pg 17 h - 3^rd^ Os 6 h. After staining, the samples were dehydrated and embedded in Spurr’s resin as described above for 2 mm samples. After resin curing, they were trimmed to expose the center, their surface was smoothed and they were imaged in high-vacuum SEM.

### UA and Ld steps for 2mm samples

Four groups of 2 mm samples (groups differ in incubation of UA and Ld steps) were stained using the following protocol: 1^st^ Os 3 h → FeCN 17 h → 2^nd^ Os 3 h → CaC 0.5 h → H_2_O 0.5 h (twice) → Pg in water 17 h) → H_2_O 0.5 h (twice) → 3^rd^ Os 6 h → H_2_O 0.5 h (twice) → 2% or 4% UA (17 h at 4°C, 2 h at 50°C) → H_2_O 0.5 h (twice) → Ld 50 °C, 4h or 24 h → H_2_O 0.5 h (twice). The combination of UA and Ld incubation were as follows: (1) 2% UA-No Ld; (2) 2% UA-Ld 4 h; (3) 4% UA-No Ld; (4) 4% UA-Ld 24 h. After staining, the samples were dehydrated and embedded in Spurr’s resin as described above for 2 mm samples. After resin curing, samples were imaged in μCT and then trimmed to expose the center, their surface was smoothed and imaged in high-vacuum SEM.

### Temperature of Os incubation step

Two groups of 2 mm samples were stained according to the following steps: (1) Os 4°C 7 days; or (2) Os 4°C 7 days, RT 1 day. After staining, the samples were dehydrated and embedded in Spurr’s resin as described above for 2 mm samples. After resin curing, they were trimmed to expose the center, their surface was smoothed and they were then imaged in low-vacuum SEM.

### Os incubation at 4 °C followed by FeCN incubation

Four groups of samples were stained according to the following steps: (1) Os 4°C 6 days, RT 1day → FeCN RT 1 day; (2) Os 4°C 6 days, RT 1 day → FeCN 4°C 1 day; (3) Os 4°C 7 days → FeCN RT 1 day; (4) Os 4°C 7 days → FeCN 4°C 1 day. After staining, the samples were dehydrated and embedded in Spurr’s resin as described above for 2 mm samples. After resin curing, they were trimmed to expose the center, their surface was smoothed and they were imaged by low-vacuum SEM.

### Diffusion of Os at 4°C in hemisphere samples

A hemisphere was incubated in OsO_4_ at 4 °C. At different time points (17 h, 24 h, 40 h), it was taken out of the fridge to perform a fast μCT scan (usually about 15-20 min total) and then returned to the fridge at 4°C. μCT images were analyzed using Zeiss TXM3DViewer software, in which the depth of OsO_4_ incubation was measured on sagittal reslices.

### Effect of CaC steps on staining gradient in hemispheres

Two groups of hemispheres were stained with the following two conditions: (1) Os 48 h → FeCN 48 h; or (2) Os 48 h → CaC 48 h → FeCN 48 h. After staining, the hemispheres were briefly rinsed in CaC and cut into coronal sections of about 2 mm thickness with a razor blade. Afterwards, the coronal sections were dehydrated and embedded in Spurr’s resin as described above for 2 mm samples. After resin curing, they were imaged in μCT, and then trimmed flat to expose the top surface, smoothed and imaged by low-vacuum SEM.

### Effect of CaC steps on velocity of FeCN diffusion

Two groups of 2 mm samples were stained with the following two conditions: (1) Os 24 h → FeCN 1.5 h; or (2) Os 24 h → CaC 24 h → FeCN 1.5 h. After staining, the samples were dehydrated and embedded in Spurr’s resin as described above for 2 mm samples. After resin curing, they were imaged in μCT, and then trimmed flat to expose the top surface, smoothed and imaged by low-vacuum SEM and EDS. For EDS measurement, we selected point measurement in the neuropile of the sample center.

### Interaction of Pg incubation with Os-FeCN gradient (H3)

A hemisphere sample (H_3_) was stained according to the following steps: 1^st^ Os 4°C 96 h, RT 24 h→CaC 4°C 48 h → FeCN 4°C 48 h → 2^nd^ Os 72 h→CaC 24 h → H_2_O 29 h → Pg 24 h → H_2_O 48 h → 3^rd^ Os 48 h → 4% UA 4°C 48 h, 50°C 5 h→H_2_O 42 h. For all CaC and H_2_O steps, the corresponding solutions were changed every 4 h/overnight (i.e. once in the morning, noon and afternoon). During staining, the hemispheres were imaged in μCT at the main staining steps (after FeCN, after 2^nd^ Os, after Pg). After staining, the hemisphere was cut into about 2 mm coronal sections with a razor blade. Afterwards, the coronal sections were dehydrated and embedded in Spurr’s resin as described above for 2 mm samples. After resin curing, they were trimmed flat to expose the top surface, smoothed and imaged by high-vacuum SEM.

### Interaction of Pg incubation with Os-FeCN gradient and CaC (H1,2)

Two hemisphere samples (H1,2) were stained according to the following steps: 1^st^ Os 4°C 63 h, RT 24 h→CaC 4°C 24 h → FeCN 4°C 48 h → CaC 24 h → 2^nd^ Os 32 h→CaC 17 h → H_2_O 9 h → Pg 48 h → H_2_O 40 h → 3^rd^ Os 48 h → 4% UA 4°C 48 h, 50°C 5 h→H_2_O 48 h. For all CaC and H_2_O steps, the corresponding solutions were changed every 4 h/overnight (i.e. once in the morning, noon and afternoon). During staining, the hemispheres were imaged in μCT at the main staining steps (after FeCN, after 2^nd^ Os, after Pg). After staining, the hemisphere was cut into about 2 mm coronal sections with a razor blade. Afterwards, the coronal sections were dehydrated and embedded in Spurr’s resin as described above for 2 mm samples. After resin curing, they were trimmed flat to expose the top surface, smoothed and imaged in high-vacuum SEM.

### Effect of CaC incubation at RT for 2 days on Os-FeCN gradient (H4,5,6)

Three hemisphere samples (H4,5,6) were stained according to the following steps: 1^st^ Os 4°C 72 h, RT 24 h→CaC 48 h → FeCN 4°C 48 h → CaC 24 h → 2^nd^ Os 48 h→CaC 24 h → H_2_O 24 h → Pg 24 h → H_2_O 24 h → 3^rd^ Os 24 h → 4% UA 4°C 48 h, 50°C 5 h→H_2_O 48 h. For all CaC and H_2_O steps, the corresponding solutions were changed every 4 h/overnight (i.e. once in the morning, noon and afternoon). During staining, the hemispheres were imaged in μCT at the main staining steps (after FeCN, after 2^nd^ Os, after Pg). After staining, H5 was cut into about 2 mm coronal sections with a razor blade. Afterwards, the coronal sections were dehydrated and embedded in Spurr’s resin as described above for 2 mm samples. H4,6 were embedded in Spurr’s resin. After resin curing, H5 was trimmed flat to expose the top surface, smoothed and imaged by high-vacuum SEM. H4,6 was trimmed to expose the center, surface smoothed and imaged by high-vacuum SEM.

### Effect of CaC incubation at 4°C for 4 days on Os-FeCN gradient (H13)

A hemisphere sample (H13) was stained according to the following steps: 1^st^ Os 4°C 68 h →CaC 4°C 96 h → FeCN 4°C 72 h → CaC 48 h → 2^nd^ Os 48 h→CaC 24 h → H_2_O 24 h → Pg 24 h → H_2_O 24 h → 3^rd^ Os 41 h → 4% UA 4°C 48 h, 50°C 12 h→H_2_O 24 h. For all CaC and H_2_O steps, the corresponding solutions were changed every 4 h/overnight (i.e. once in the morning, noon and afternoon). During staining, the hemispheres were imaged in μCT at the main staining steps (after 2^nd^ Os, after Pg). After staining, the hemisphere was embedded according to Spurr’s resin embedding for hemispheres. After resin curing, it was trimmed to expose the center, surface smoothed and imaged by high-vacuum SEM.

### Whole brain staining (W1)

A mouse whole brain sample (W1) was stained according to the following steps: 1^st^ Os 4°C 96 h →CaC 4°C 168 h → FeCN 4°C 72 h → CaC 4°C 48 h, RT 48 h → 2^nd^ Os 48 h→CaC 72 h → H_2_O 48 h → Pg 24 h → H_2_O 48 h → 3^rd^ Os 96 h → 4% UA 4°C 48 h, 50°C 5.5 h→H_2_O 24 h. For all CaC and H_2_O steps, the corresponding solutions were changed every 4 h/overnight (i.e. once in the morning, noon and afternoon). During staining, the brain was imaged in μCT at the main staining steps (during 1^st^ Os, after 2^nd^ Os, after Pg). After staining, the brain was cut into ~2 mm thick coronal sections. Then the sections were embedded according to Spurr’s resin embedding for 2 mm samples. After resin curing, the sections were trimmed flat to expose the surface, smoothed and checked by high-vacuum SEM.

### Effect of 2^nd^ Os step on sample stability in H_2_O

Two groups of 2 mm samples (the groups differed in whether they were exposed to a 2^nd^ Os step) were incubated in the following conditions: (1) 1^st^ Os 24 h →CaC 24 h → FeCN 24 h →2^nd^ Os 3 h →CaC 0.5 h → H_2_O (two times 0.5 h) → H_2_O; or (2) 1^st^ Os 24 h →CaC 24 h → FeCN 24 h →CaC 0.5 h → H_2_O (two times 0.5 h) → H_2_O. Afterwards, the samples were kept in H_2_O, and imaged in the light microscope at different time points (from 2 h to 100 h) to investigate macroscopic sample integrity.

### Dehydration of 2 mm samples

After the last water rinsing step in the staining protocol, 2 mm samples were exposed to graded dehydration series in ethanol (50% 4°C, 75% 4°C and 100% RT; each step 45 min) and then three rounds of acetone incubation (each 45 min).

### Dehydration of hemispheres

After the last water rinsing step in the staining protocol, hemisphere samples were exposed to graded dehydration series in ethanol (25% 4°C, 50% 4°C, 75% 4°C and 100% RT; each step 8 h/overnight) and then three rounds of acetone incubation (each time 8 h/overnight).

### Infiltration of Spurr’s resin and embedding of 2 mm samples

After dehydration, a graded incubation of Spurr’s resin (0.95 g ERL 4221, 5.9 g DER 736, 0.1 g NSA, 113 ul DMAE) was applied at 25%, 50%, 75% in acetone (each step 8 h or overnight). Afterwards, the samples were incubated for 4 rounds in 100% Spurr’s resin (each time 8 h or overnight). Finally, they were embedded in freshly prepared Spurr’s resin and cured at 70°C for 1-3 days. All steps were performed at 4°C. For each resin exchange, the tubes were taken out from fridge 20-30 min before to be warmed to RT.

### Infiltration of Spurr’s resin and embedding of hemisphere samples

After dehydration, a graded incubation of Spurr’s resin (0.95 g ERL 4221, 5.9 g DER 736, 0.1 g NSA, 113 ul DMAE) was applied at 25%, 50%, 75% in acetone (each step 24 h). Afterwards, the samples were incubated for 2 days in 90% resin, for 3 days in 95% resin and for 3 days 100% resin (change every 8 h or overnight for 90%, 95% and 100% resin steps). Finally, they were embedded in freshly prepared Spurr’s resin and cured at 70°C for 1-3 days. All resin steps were performed at 4°C. For each resin exchange, the tubes were taken out from fridge 20-30 min before to be warmed to RT.

### Measurement of viscosity of Epon resin

10 ml Epon resin (Sigma-Aldrich, with a ratio of 5.9 g Epon medium, 2.25 g DDSA, 3.7 g MNA, and 205 ul DMP) was prepared and stored in 15 ml Falcon tubes at either 4 °C or RT. At different time points, videos were acquired of the resin moving in the tube after been turned up-side down. The distance of movement of the resin in the tube was measured using the ticks (ml) printed on the Falcon tubes. The speed of the resin movement was used as a measurement of the resin’s viscosity.

### Infiltration with Epon resin and embedding of 2 mm samples

After dehydration, samples were incubated in pure acetone for 3 times, each time 45 min. Then a graded incubation of Epon resin (5.9 g Epon medium, 2.25 g DDSA, 3.7 g MNA, 205 ul DMP) was applied at 12.5%, 25%, 37.5%, 50%, 62.5%, 75% to 87.5% in acetone (each step 4 h or overnight). Afterwards, the samples were incubated for 4-6 rounds in 95% Epon resin (each time 8 h or overnight), and then for 4 rounds in 100% Epon resin (each time 8 h or overnight) before embedding for curing at 60 °C for 1-3 days. All resin steps were carried out at 4°C. For each resin exchange, the tubes were taken out from fridge 20-30 min before for warming to RT.

### Light-microscopic imaging of sample surface to assess resin gradient

Resin embedded samples (2-3 mm/hemispheres) were trimmed to expose the center and surface smoothed with a diamond knife. Light-microscopic images of the sample surface were acquired with a slight tilt of the imaging plane, such that the otherwise black sample surface appeared as silver-colored.

### μCT volumetric imaging of samples without resin embedding

To allow imaging in μCT without the need for resin embedding, 2 mm samples were embedded instead in 2% agarose (Sigma-Aldrich) in 0.15 M CaC or in water (depending on the last staining step that the sample was exposed to) in 2 ml Eppendorf tubes. They were imaged in μCT (Zeiss Xradia 520 Versa) using a voltage of 80 kV at voxel size of 3-6 μm. For μCT imaging of hemispheres, the samples were kept in the 50 ml glass tubes. Then, the glass tubes were put into a 140 ml syringe to be kept stable during μCT imaging (using voxel size of 10-60 μm).

### Low-vacuum SEM imaging of incompletely stained samples

To investigate samples at intermediate staining steps without complete staining (and therefore often reduced signal and conductivity), these were resin embedded. Then they were trimmed with a diamond head trimmer (Leica EM TRIM2) to expose the center to the block faces of samples; the block face was smoothed with a diamond knife ultra-microtome (Leica EM UC7); and then imaged in a scanning electron microscope with a field-emission cathode and low vacuum mode (Quanta FEG 450, FEI Company). The chamber pressure was set to 30 Pa. For the incident electron beam, a spot size of 3.5 and acceleration energy of 5 keV were used for imaging at a pixel dwell time of 8-20 μs and a pixel size of 5.62 nm^2^ in-plane (corresponding electron dose: 112-280 e-/nm^2^ without considering the electron loss caused by the skirting effect of low-vacuum) or 11.24 nm^2^ in-plane (corresponding electron dose: 28-70 e^−^/nm^2^, without considering the electron loss caused by the skirting effect of low-vacuum), at working distance of about 5 mm using the back scattered electron CBS detector.

### High vacuum SEM imaging of completely stained samples

For those samples that were fully stained and resin embedded (and therefore were expected to show sufficient conductivity), trimming and smoothing was similar to the previous section, but SEM imaging was performed in high-vacuum mode (5*10^−4^ Pa). For the incident electron beam, a spot size of 3.5 and acceleration energy of 2.8 keV was used for imaging at a pixel dwell time of 6-8 μs and a pixel size of 11.24 nm^2^ in-plane (corresponding electron dose: 16-21 e^−^/nm^2^) or 5.62 nm^2^ (corresponding electron dose: 64-84 e^−^/nm^2^), at a working distance of about 5 mm using the back scattered electron CBS detector.

### EDS analysis

Resin embedded samples were trimmed to expose the center as a block face (Leica EM TRIM2), smoothed (Leica EM UC7), and coated with a 10 nm gold layer (Leica EM ACE600); afterwards, they were imaged in a scanning electron microscope (Amray 1830) equipped with an Si(Li) EDS detector. An incident electron beam with energy of 18 keV was used at working distance of 15-20 mm and a takeoff angle of 20.4 degree. The spectrum collection time was 20-90 s.

### UV-vis spectrum acquisition of Os(vi) solution

We prepared 1% potassium osmate(vi) in 0.15 M sodium cacodylate buffer by adding 0.05 g potassium osmate(vi) powder (Sigma-Aldrich) into 5 ml cacodylate buffer. The solution was diluted 100 times and put into a glass cuvette to avoid spectrum signal clipping. The measurement was performed using the wavelength range 190-1400 nm on a UV-vis spectrometer (Jasco V-670). Measurements on the same solution were made at time points 2 min, 2 h and 24 h.

### Raman spectrum measurement of Os vi + FeCN solutions

We measured the following chemicals without dilution in a custom built Raman spectrometer (Chemistry department of Goethe University Frankfurt) with a range of 0-4,400 cm^−1^ wavelength: (1) 0.15 M sodium cacodylate buffer; (2) 1% potassium osmate(vi) in 0.15 M cacodylate buffer; (3) 0.3% potassium osmate(vi) and 1.3% potassium ferrocyanide(ii) (Sigma Aldrich) in 0.15 M cacodylate buffer; (4) 2.5% potassium ferrocyanide(ii) in 0.15 M cacodylate buffer; (5) 1.9% potassium ferricyanide(iii) (Sigma-Adrich) in 0.15 M cacodylate buffer; (6) Staining solution of 2 mm samples in 2 ml Eppendorf tubes, after 24 h of 2% OsO4 (Serva) in 0.15 M cacodylate buffer, 1 h 0.15 M cacodylate buffer wash, and 17 h of 2.5% potassium ferrocyanide(ii) in 0.15 M cacodylate buffer.

