## Supplementary Material for "High-contrast en-bloc staining of mouse whole-brain samples for EM-based connectomics"

### Supplementary Results

#### Chemical concepts of $OsO_4$-FeCN-related enhancement of membrane contrast

For enhanced membrane contrast of biological specimens in electron microscopy, potassium hexacyanoferrate(ii) (FeCN) has been combined with $OsO_4$ since the 1970s (Bruijn, 1973; Karnovsky, 1971). Chemically, this was considered to involve a redox reaction between $OsO_4$ and FeCN(Bruijn, 1973; Karnovsky, 1971; Riemersma, Alsbach, & Bruijn, 1984; White, Mazurkiewicz, & Barrnett, 1979), therefore the protocol has been referred to as “reduced-osmium” (rO) protocol. Two mechanisms were considered for the contrast enhancement in the rO protocol: Either more osmium compounds were deposited into membranes(Bruijn, 1973; Bruijn & Breejen, 1975; White et al., 1979; Willingham & Rutherford, 1984), or osmium compounds in the cytosolic background were removed(Litman & Barrnett, 1972; Neiss, 1984; Willingham & Rutherford, 1984). A more recent improved staining protocol was based on the following chemical logic(Hua, Laserstein, & Helmstaedter, 2015): The staining process was considered as a two-step chemical reaction: First, FeCN reduces $Os^{viii}O_4$ to its lower valence form : $Os^{vi}O_2(OH)_4^{2-}$ (Os(vi)). Then, two molecules of Os(vi) dismutate into one molecule of $OsO_4$ and one molecule of $Os^{IV}O_2$, and $OsO_2$ was assumed to be able to deposit into membranes(Hua et al., 2015), thus result in higher membrane contrast.

For the protocols reported here, we developed a different chemical concept. First, we noted that several experimental results would not support the above “redox-dismutation” (Hua et al., 2015) explanation. The assumed dismutation reaction was found in either organic solvents or low pH water solution(Korn, 1967), but it is less likely to occur in cacodylate buffer (CaC). In support of this, we found no change in UV-Vis absorption spectrum of potassium osmate (vi) ($K_2[Os^{VI}O_2(OH)_4]$) solution in CaC after 24 h compared to freshly prepared solution (Supplementary Fig.5d). In brain tissue, dismutation might still occur in a more complex way involving interactions with lipid membranes. However, we found that membrane contrast was not enhanced by staining samples directly with potassium osmate(vi) in CaC solution (Supplementary Fig.5c). Furthermore, we noticed that the redox reaction between $OsO_4$ and FeCN as the basis for membrane contrast enhancing(Bruijn, 1973; Hua et al., 2015; Karnovsky, 1971; Riemersma et al., 1984; White et al., 1979) had little direct support. A direct test would be to determine membrane contrast with a suppressed redox reaction. When adding an additional CaC washing step to remove $OsO_4$ from the tissue before adding FeCN, in an attempt to suppress $OsO_4$-FeCN redox reactions, similar membrane contrast was observed (Supplementary Fig.5a,b, comparing to Fig.1d,e); indicating that the $OsO_4$-FeCN redox reaction may not be a prerequisite for enhanced membrane contrast.

For developing an alternative explanation of membrane contrast enhancement, we were guided by the observation that the duration of the $OsO_4$ incubation step was critical for the effect of FeCN: when the $OsO_4$ incubation duration was short (but sufficient to stain a 2 mm-sized sample homogeneously, e.g. 3 hours), addition of FeCN actually did not enhance membrane contrast (Supplementary Fig.2a); only when prolonging the $OsO_4$ incubation to 24 h, the subsequent FeCN step was effective in membrane contrast enhancement (Supplementary Fig.2b). Which processes could have occurred during the prolonged $OsO_4$ incubation? After short incubation, both the membranes and background appeared stained (Supplementary Fig.2a), implying that osmium compounds were deposited/attached at both targets. After prolonged incubation, however, the cytosolic background (which may mainly be constituted by proteins), may have been altered by $OsO_4$ such that FeCN can react to yield products that can be washed out, reducing background staining (Supplementary Fig.2b; consistent with insights in earlier studies(Litman & Barrnett, 1972; Neiss, 1984; Willingham & Rutherford, 1984)). One possible process/reaction that may have happened between $OsO_4$ and background is protein over-oxidization(Hayat, 1981), which might induce 3D -conformational changes of the proteins (e.g. de-gelation(Porter & Kallman, 1953)). As a result, more protein-bound osmium (of valence vi(Hayat, 1981)) could be exposed to the aqueous phase, enabling the interaction with FeCN in the aqueous solution. This notion of conformational instability of proteins as a prerequisite for FeCN effects was supported by an experiment in which we added $CaCl_2$ as a protein crosslinking cation(Hayat, 1981), and indeed found this to inhibit the membrane contrast enhancement effect of FeCN after prolonged $OsO_4$ incubation (Supplementary Fig. 5g). In addition, when incubating at 4 °C, protein conformation would be expected more stable, and chemical reactions slowed down. In fact, low-temperature incubation could also inhibit the effect of FeCN on membrane contrast after prolonged $OsO_4$ incubation (Supplementary Fig.5h).

But is the addition of FeCN a necessity after prolonged $OsO_4$ exposure for enhancement of membrane contrast, or could sufficiently long OsO4 incubation alone provide sufficient membrane contrast? We stained samples with further extended $OsO_4$ incubation (from 24 h to 3-6 days) and investigated membrane contrast afterwards in low-vacuum SEM. After 6 days of $OsO_4$ incubation, enhanced membrane contrast was in fact achieved (Supplementary Fig.2c,d). However, extensive background extractions also occurred (Supplementary Fig.2c,d). Together, we could interpret this as an indication that FeCN was not strictly required for enhanced membrane contrast; however it could accelerate contrast enhancement such that sufficient contrast could be achieved before the $OsO_4$ incubation yielded extraction of background ("over-fixation").

We still needed to form a hypothesis for the actual reactions occurring between Os-compounds and FeCN. These experiments and concepts are described in the following.

During prolonged $OsO_4$ incubation, we noticed a pink reaction product diffusing out from the samples (for comparison, $OsO_4$ was yellowish in solution) (Supplementary Fig.6a), the typical color of osmate(vi) in cacodylate. This further supported the previous notion that the background osmium species exposed by prolonged $OsO_4$ was Os(vi). During the very long $OsO_4$ incubation, as the proteins were further oxidized, the exposed Os(vi) might be released into aqueous phase, likely coordinated with the anion from the CaC buffer as this could stabilize Os(vi) in water (Supplementary Fig.5d)). In this coordinated form, Os(vi) could diffuse out of the sample, reducing background staining, thus enhancing membrane contrast. This implied that this Os(vi)-removal process would be slow, as 24 h $OsO_4$ incubation had not yielded enhanced membrane contrast, but only 3-6 days of $OsO_4$ incubation (Supplementary Fig.2c,d) - consistent with the notion from previous studies that “reduced osmium” diffusion was slow(Hua et al., 2015; Mikula & Denk, 2015).

How could the addition of FeCN accelerate this Os(vi) background removal process? After $OsO_4$ incubation we would expect Os(viii) and Os(vi) freely moving in the aqueous phase. Since we consider redox reaction or dismutation unlikely as key processes in our explanation (see above), thus excluded the possibility of reaction between Os (viii)-FeCN or Os(vi)-Os(vi); we had to considered the remaining likely chemical reaction to occur between Os(vi) and FeCN. This would be consistent with the observation that the staining solution after the FeCN step had a green-blue color, similar to the color observed in the in-vitro reaction of Os(vi) and FeCN (Supplementary Fig.6a).

What would be a realistic reaction between Os(vi) and FeCN to occur? We explored this by measuring the Raman spectrum of the mixture solution of Os(vi) and FeCN. In the mixture, we did not find the spectral peak expected for potassium hexacyanoferratte (iii) (FeCNiii) (Supplementary Fig.6b), indicating that Os(vi) was not able to oxidize FeCN to FeCNiii. On the other hand, Os(vi) was known to have the capacity to coordinate with ligands like $OH^-$ (Collinson & Schroeder, 2011; Hua et al., 2015) and $CN^-$(Collinson & Schroeder, 2011; Griffith, 1962; Frantisek Opekar & Premysll Beran, 1977; František Opekar & Přemysl Beran, 1977), and FeCN was known to be unstable such that it would dissociate to yield free $CN^-$ spontaneously at low rate in aqueous solution(Tirler, Persson, Hofer, & Rode, 2015). Thus a possible reaction between Os(vi) and FeCN was a coordination reaction, in which Os(vi) competed for the $CN^-$ from FeCN and formed a coordination complex.

This hypothesis was further supported by our measurement of the Raman spectroscopy: we found a shift of the vibration peak of $O=Os^{VI}=O$ in the Os(vi)-FeCN solution compared to Os(vi) alone (Supplementary Fig.6b), indicating a ligand change in the Os(vi) molecules. The same shifted peak was also found in the brain staining solution after the FeCN step (Supplementary Fig.6c), indicating that FeCN in the sample staining processes could in fact react with Os(vi).

Together, a possible function of FeCN is to “carry” $CN^-$ to diffuse through the sample and donate $CN^-$ to exposed Os(vi) (after prolonged $OsO_4$ incubation). If the incubation of

$OsO_4$ before FeCN was not sufficiently long to expose background Os(vi), FeCN incubation would only remove the free Os(vi) in solution, not the Os(vi) bound to cytosolic proteins, thus would not result in enhanced membrane contrast. If the incubation of $OsO_4$ was sufficiently long to expose protein-bound Os(vi), FeCN incubation would however coordinate with the exposed protein-bound Os(vi), enabling its removal from the sample, and yielding enhanced membrane contrast. The likely form of this assumed complex might be $OsO_2(OH)_2(CN)_2^{2-}$, $OsO_2(CN)_4^{2}$-(Frantisek Opekar & Premysll Beran, 1977; František Opekar & Přemysl Beran, 1977).

This concept adds a new view on the long mysterious chemical nature of $OsO_4$-FeCN membrane contrast enhancement (Supplementary Fig.6d): (1) $OsO_4$ stained both membrane and cytosolic proteins (background) during short incubation; (2) With prolonged incubation, $OsO_4$ over-oxidized the background proteins and thereby exposed protein-bound Os(vi) to aqueous phase; (3) FeCN would carry $CN^-$ through the sample, and provide $CN^-$ to the exposed protein-bound Os(vi) to form a stable coordination compound (note this would be a compound between Os(vi) and $CN^-$, but not between Os(vi) and FeCN itself as proposed in 1970s(White et al., 1979)). The coordination compound between Os(vi) and $CN^-$ could then easily diffuse out(Bruijn & Breejen, 1975).

**Discussion of previous protocols in the framework of the exposure-coordination based staining concept**

The possible exposure-coordination concept of membrane contrast enhancement would add a surprising view on Karnovsky(Karnovsky, 1971) and de Bruijn's(Bruijn, 1973) discovery in 1970s to find FeCN as a staining enhancement agent: the reason this would have worked was that $Fe^{2+}$ was strong enough to maintain $CN^-$ in the coordination complex, such that $CN^-$ would not react directly with the osmium compounds deposited within membranes (as the osmium compounds in membranes are those we would like to keep). In fact, otherwise the effect would be dramatic: the staining of the entire sample would be lost if osmicated samples were incubated in KCN which provide free $CN^-$(Bruijn & Breejen, 1975). FeCN, in contrast, would be more strongly coordinated and only be able to donate $CN^-$ when encountering the exposed/free Os(vi) in the aqueous solution, forming the stable coordination product of Os(vi) and $CN^-$ (Frantisek Opekar & Premysll

Beran, 1977; František Opekar & Přemysl Beran, 1977) that would be easily washed out(Bruijn & Breejen, 1975). Our chemical concept was also consistent with the “background washing” insight from the early days(Litman & Barrnett, 1972; Neiss, 1984; Willingham & Rutherford, 1984).

The same incubation time dependent effect would also exists in the original rO protocol (Supplementary Fig.5e,f). The principles summarized above can be applied similarly: even though FeCN and $OsO_4$ were mixed together in rO (which would yield a mixture of $OsO_4$, FeCN, FeCNiii and Os vi); however, FeCN-based reactions would still only be occurring once the background proteins were exposing Os(vi) for $CN^-$ - complex formation.

To provide a possible explanation for the staining gradient and precipitation band that would appear when staining samples with rO that were larger than ~200 µm in size(Hua et al., 2015; Mikula & Denk, 2015), we considered the following: The very slow centripetal diffusion of Os(vi) (Supplementary Fig.5c) from all sides of the sample could yield a density of Os(vi) at a certain depth such that CaC could no longer stabilize the high amount of Os(vi), and local dismutation would occur (thereby yielding $Os^{IV}O_2$, Supplementary Fig. 5c,d).

To evaluate the rO(Bruijn, 1973; Karnovsky, 1971), Hua(Hua et al., 2015) and Mikula(Mikula & Denk, 2015) protocols in the framework of the exposure-coordination concept, we consider the Os(vi) coordination reactions with different ligands (hydroxide, cacodylate, cyanide and others, Supplementary Fig.6e). The coordination bonds from different ligands would differ in their strength, influencing the stability of the complexes. Similarly, the complexes would also differ in size as a result of different ligand sizes; this would influence the mobility of the complexes during diffusion. In water, Os(vi) would coordinate with hydroxide ions ($OH^-$) and form an unstable complex which would dismutate relatively easily(Hua et al., 2015; Korn, 1967). In cacodylate buffer, Os(vi) would form more stable complexes with cacodylate anions ($(CH_3)_2AsO_2^-$), very likely also by coordination bond(Collinson & Schroeder, 2011), which should be stronger so that it can inhibit Os(vi) dismutation (Supplementary Fig.5d). The difference in the size of the coordination bonds by hydroxide vs. cacodylate could also contribute: in hydroxide coordination there would be a distance of only two chemical bonds (Os-O-H)), but in cacodylate coordination there would be a distance of four chemical bonds (Os-O-As-C-H), potentially affecting final compound size and mobility, possibly explaining why the [Os(vi)-cacodylate] coordination compound diffused slowly(Hua et al., 2015; Mikula & Denk, 2015).

For the formamide(Mikula & Denk, 2015) used in Mikula's protocol, Os(vi) would coordinate with the amine group in formamide(Collinson & Schroeder, 2011). The formamide coordination with Os(vi) would result in a three bond distance (Os-N-C-H), thus less than the four bond distance of cacodylate; this might give the formamide coordination compound more mobility. The coordination ligand amine group should also form relatively stronger coordination than hydroxide(Collinson & Schroeder, 2011), this could help preventing precipitations as discussed in the "reduced osmium" protocol.

**Remaining artifacts**

Three types of artifacts were observed in the large-sample protocols: (1) Blood vessels partly detached from the surrounding neuropil (Supplementary Fig.7b); (2) Remaining micro-breakages in subcortical neuropil (Supplementary Fig.7d,e); (3) Damage to outer parts of the cerebellum that could break off during water incubation steps (Supplementary Fig.7f).

For blood vessel artifacts, in order to determine in which protocol step they appeared, we imaged 2 mm samples after 24 h RT $OsO_4$ incubation in μCT; with this method, we avoided the influence of dehydration and resin infiltration. Blood vessels already detached after the first $OsO_4$ incubation step (Supplementary Fig.7c). The result was the same for samples that was $OsO_4$ incubated at 4°C for 4 days (Supplementary Fig.7c). While not a major concern for connectomics projects aimed at synaptic circuitry, for projects in which the complete integrity of blood vessels were to be essential, we see the need for further targeted optimization of the protocol.

For micro-breakages in subcortical areas, these were rather small (≤ 500 nm in width, see in Supplementary Fig.7d), rendering detection by μCT difficult. Since these occurred rarely and predominantly in myelin-rich areas, we expect neurite reconstruction to be largely unaffected.

For the cerebellum breakage artifacts, it was obvious from the staining process that they occured during the first $H_2O$ incubation step (Supplementary Fig.7g). We found this to be influenced by the temperature of CaC incubation between $OsO_4$ and FeCN: when the CaC incubation step was at 4°C, the cerebellum was preserved much better than at RT (Supplementary Fig.7g).

For stabilizing the samples in water, we found a second $OsO_4$ step after FeCN to be useful (Supplementary Fig.3i). Another approach would be to increase the osmolarity of the solution with some inactive chemicals. However, by our experience, to find a chemical to balance the osmolarity of water before and after pyrogallol would be a difficult task. We have observed that CaC buffer won't work as it would cause gradients and breakages (Supplementary Fig.3g), which we think is caused by the coordination interaction between CaC and Os(vi). However, Os(vi) is very active in coordination chemistry(Collinson & Schroeder, 2011). Thus it would be another chemical screening project to find out a chemical that does not coordinate with osmium species to be used as osmolarity balancer.

**Supplementary Tables**

**Supplementary Table 1: Detailed protocols for 2-3 mm and mouse hemisphere/whole brain sample staining and resin embedding**Samples sized 2-3 mm were stained in 2 ml Eppendorf tubes, hemispheres / whole brains were stained in 50 ml tubes. All steps were at room temperature (RT) unless otherwise noted. From step 3 (FeCN) onwards, all tubes were covered with aluminum foil.

| **I. Staining** | | | | | | |
|---|---|---|---|---|---|---|
| **Step** | **Solution** | **Conc.** | **2-3 mm** | **Hemisphere (H)** | **Whole Brain (WB)** | **Notes** |
| **1a** | **$OsO_4$** | 2% in 0.15 M CaC, pH 7.4 | n/a | 72 h, 4°C | 96 h, 4°C | |
| **1b** | **$OsO_4$** | 2% in 0.15 M CaC, pH 7.4 | 24 h | n/a | n/a | |
| **2** | **CaC** | 0.15 M, pH 7.4 | 1.5-2 h, 4°C<br><br>change every 0.5 h | 96 h, 4°C<br><br>change every 4 h/o.n. | 168 h, 4°C<br><br>change every 4 h/o.n. | |
| **3** | **FeCN** | 2.5% in 0.15 M CaC, pH 7.4 | overnight (17 h) , 4°C | 72 h, 4°C<br><br>change every 24 h | 72 h, 4°C<br><br>change every 24 h | Light sensitive. H4-6 used 48h, but 72h recommended |
| **4** | **CaC** | 0.15 M, pH 7.4 | 1.5-2 h<br><br>change every 0.5 h | 48 h<br><br>change every 4 h/o.n. | 96 h<br><br>change every 4 h/o.n. | |
| **5** | **$OsO_4$** | 2% in 0.15 M CaC, pH 7.4 | 3 h | 48 h | 48 h | |
| **6** | **CaC** | 0.15 M, pH 7.4 | 0.5-1 h<br><br>change every 0.5 h | 24 h<br><br>change every 4 h/o.n. | 72 h<br><br>change every 4 h/o.n. | |
| **7** | **$H_2O$** | | 1-2 h<br><br>change every 0.5 h | 24 h<br><br>change every 4 h/o.n. | 48 h<br><br>change every 4 h/o.n. | |
| **8** | **Pyrogall ol** | 4% in $H_2O$ | overnight (17 h)<br><br>change every | 24 h | 24 h | Freshly prepared; light sensitive |

| | | | | | | |
|---|---|---|---|---|---|---|
| | | | 4h/o.n. | | | |
| **9** | **$H_2O$** | | 1-2 h<br><br>change every 0.5 h | 24 h<br><br>change every 4 h/o.n. | 48 h<br><br>change every 4 h/o.n. | |
| **10** | **$OsO_{4,H2O}$** | 2% in $H_2O$ | 6 h | 48 h | 96 h | Was 24h for H4-6, but 48h (H13) recommended |
| **11** | **$H_2O$** | | 1 h<br><br>change every 0.5 h | 24 h<br><br>change every 4 h/o.n. | 48 h<br><br>change every 4 h/o.n. | |
| **12a** | **Uranyl Acetate** | 4% in $H_2O$ | overnight (17 h), 4°C | 48 h, 4°C | 48 h, 4°C | Light sensitive. UA solution was not changed for steps 12a and 12b, only the temperature was changed. |
| **12b** | **Uranyl Acetate** | 4% in $H_2O$ | 2 h, 50°C | 5 h, 50°C | 5 h, 50°C | |
| **13** | **$H_2O$** | | 1 h<br><br>change every 0.5 h | 24 h<br><br>change every 4 h/o.n. | 48 h<br><br>change every 4 h/o.n. | |

**II. Dehydration**

| Step | Solution | 2-3 mm | hemisphere | Notes |
|---|---|---|---|---|
| **1** | **25% Ethanol in $H_2O$** | n/a | 8 h/o.n., 4°C | |
| **2** | **50% Ethanol in $H_2O$** | 0.5 h, 4°C | 8 h/ o.n. 4°C | |
| **3** | **75% Ethanol in $H_2O$** | 0.5 h, 4°C | 8 h/ o.n., 4°C | |
| **4** | **Pure Ethanol** | 45 min, RT | 8 h/ o.n.., RT | Samples can stay in pure Ethanol or pure acetone for days (up to 1 week), if a pause is needed practically |
| **5** | **Pure Acetone** | 2 h 15 min<br><br>change every 45 min | 32 h, RT<br><br>change every 8 h/o.n. | |

**III. Resin infiltration & Embedding**

**Spurr's resin**
4.1 g ERL 4221, 0.95 g DER 736, 5.9 g NSA, 113 µl DMAE

**Epon resin**
5.9 g Epon embedding medium, 2.25 g DDSA, 3.7 g MNA, 205 µl DMP

| Step | Conc. (in acetone) | 2-3 mm | Hemisphere /WB | Step | Conc. (in acetone) | 2-3 mm |
|---|---|---|---|---|---|---|
| **1** | **25% resin** | 8 h/o.n., 4°C | 24 h, 4°C | **1** | **12.5% resin** | 4 h/o.n., 4°C |

| | | | | | | |
|---|---|---|---|---|---|---|
| **2** | **50% resin** | 8 h/o.n., 4°C | 24 h, 4°C | **2** | **25% resin** | 4 h/o.n., 4°C |
| **3** | **75% resin** | 8 h/o.n., 4°C | 24 h, 4°C | **3** | **37.5% resin** | 4 h/o.n., 4°C |
| **4** | **90% resin** | n/a | 2 days, 4°C,<br><br>change every 8h/o.n. | **4** | **50% resin** | 4 h/o.n., 4°C |
| **5** | **95% resin** | n/a | 3 days, 4°C,<br><br>change every 8h/o.n. | **5** | **62.5% resin** | 8 h/o.n., 4°C |
| **6** | **100% resin** | 2 days, 4°C;<br><br>change every 8h/o.n. | 3 days, 4°C,<br><br>change every 8h/o.n. | **6** | **75% resin** | 8 h/o.n., 4°C |
| **7** | **embed** | 70°C, 3 days | | **7** | **87.5% resin** | 8 h/ o.n., 4°C |
| For all resin steps at 4°C, the tubes were taken out of fridge for 30 min (cap closed) before change to the solution of the next step to warm up to room temperature to avoid moisture. | | | | **8** | **95% resin** | 3 days, 4°C<br><br>change every 8h/o.n. |
| | | | | **9** | **100% resin** | 2 days, 4°C;<br><br>change every 8h/o.n. |
| | | | | **10** | **embed** | 60°C, 3 days |

For all solution changing step, we tried to remove as much old solution as possible to make sure there was as little interaction with new solutions as possible. For samples with sizes in between the listed 2-3 mm and hemisphere and whole brain volumes, interpolation of the above steps should be applied, and 2 ml Eppendorf tubes should also be changed to larger volume tubes. If modifications were made based on our recommended protocol for intermediately sized volumes, we also recommend checking staining homogeneity in μCT and SEM after the major staining steps (i.e., 1$^{st}$ $OsO_4$, FeCN, 2$^{nd}$ $OsO_4$, Pg, 3$^{rd}$ $OsO_4$, UA) to verify the validity of any modifications.

**Supplementary Table 2. Employed chemicals**

| Chemical | Name | Preparation |
|---|---|---|
| MilliQ water | $H_2O$ | - |
| 0.15 M sodium cacodylate buffer | CaC | Diluting 0.3 M CaC (Sigma-Aldrich) 1:1 with water, pH adjusted to 7.4 by 1M NaOH (Sigma-Aldrich) |
| 2% $OsO_4$ in 0.15 M cacodylate buffer | Os | Diluting 4% $OsO_4$ (Serva) 1:1 with 0.3 M CaC (pH 7.4), no further pH adjustment |
| 2% $OsO_4$ in water | Os-aqua | Diluting 4% $OsO_4$ (Serva) 1:1 with water |
| 2.5% potassium ferrocyanide in 0.15 M cacodylate buffer | FeCN | Dissolving 1.25 g FeCN (Sigma-Aldrich) in 50 ml 0.15 M CaC (Sigma-Aldrich) |
| 1% $OsO_4$ + 1.5% FeCN in 0.15 M CaC | rO | Diluting 2% $OsO_4$ (Serva) in 0.15 M CaC (Sigma-Aldrich) 1:1 with 2.5% FeCN (Sigma-Aldrich) in 0.15 M CaC (Sigma-Aldrich) |
| 1% thiocarbohydrazide in water | TCH | Dissolving 0.5 g TCH (Sigma-Aldrich) in 50 ml water, shake for 1 h, filter before use. Light sensitive, cover with aluminum foil |
| 3.75% pyrogallol in water | Pg | Dissolving 1.87 g Pg (Sigma-Aldrich) in water |
| 3.75% pyrogallol in CaC | Pg-CaC | Dissolving 1.87 g Pg (Sigma-Aldrich) in 0.15 M CaC (Sigma-Aldrich) |
| 2% uranium acetate in water | 2% UA | Dissolving 1 g UA (Serva) in 50 ml water, shake for 1 h, filter before use. Light sensitive, cover with aluminum foil |
| 4% uranium acetate in water | 4% UA | Dissolving 2 g UA (Serva) in 50 ml water, shake for 1 h, filter before use. Light sensitive, cover with aluminum foil |
| 0.66% lead aspartate | Ld | Dissolving 0.33 g lead nitrate (Sigma-Aldrich) in 50 ml 0.03 M aspartate buffer (pH 3.8, Serva), adjust pH to 5.0 with 1 M KOH (Sigma-Aldrich). Light sensitive, cover with aluminum foil. |

**Supplementary Table 3. Protocols for all staining experiments**

Abbreviated protocol descriptions for all shown control and main experiments with corresponding figure panels. Protocol steps were denoted as chemical, incubation time, temperature. Incubation time in h unless specified otherwise. Temperature only noted when other than room temperature (RT). Sample screening method is indicated (µCT and/or SEM); Sample size, dehydration, infiltration and embedding steps are reported in main methods. x2: applied twice.

| Exp ID | Experiment | Staining Steps (brief) | Figures |
|---|---|---|---|
| 1 | Staining 2 mm samples with 1 mm protocol | Os 1.5→FeCN 1.5→ Os 45min→ CaC 0.5→x2 $H_2O$ 0.5→ TCH 1→ x2 $H_2O$ 0.5→ Os-aqua1.5→ x2 $H_2O$ 0.5→2% UA overnight 4°C, 2 h 50°C→ µCT, SEM | Fig.1c; Suppl.Fig.1 a,b |
| 2 | Step-by-step diagnosis of 1 mm protocol | Os 1.5 (µCT)→FeCN 1.5 (µCT)→ Os 45min (µCT)→ CaC 0.5→x2 $H_2O$ 0.5 → TCH 1→ x2 $H_2O$ 0.5→ Os-aqua 1.5 (µCT)→ x2 $H_2O$ 0.5→2% UA overnight 4°C, 2 h 50 °C (µCT) | Suppl.Fig.1 c |
| 3 | Extending FeCN incubation for 2 mm | Os 3→FeCN 1.5/3/7/12/17→ Os 3→ µCT | Suppl.Fig.1 d |
| 4 | Os short vs. long on FeCN's effect | Os 3→ SEM<br>Os 3→ FeCN 17→SEM<br>Os 24→SEM<br>Os 24→ FeCN 17→ SEM | Suppl.Fig.2 a,b |
| 5 | Extending TCH incubation for 2 mm | Os 3→FeCN 17→ Os 3→ CaC 0.5→ x2 $H_2O$ 0.5 → TCH 1.5/3/5→ x2 $H_2O$ 0.5→ Os-aqua 3→ µCT | Suppl.Fig.1 e |
| 6 | Replacing TCH by Pg | Os 1.5→FeCN 1.5→ Os 45min→ CaC 0.5→x2 $H_2O$ 0.5→ TCH/$H_2O$/Pyrogallol 1→ x2 $H_2O$ 0.5→ Os acq 1.5→ x2 $H_2O$ 0.5→ 2% UA overnight 4°C, 2 h 50°C→ SEM & EDS | Suppl.Fig.1f ,g,h |
| 7 | Comparing Pg in H2O vs. CaC | Os 24→FeCN 17→ Os 3→ CaC 0.5→x2 $H_2O$ 0.5→ Pg-aqua/Pg-CaC 17→ x2 $H_2O$ 0.5→ Os-aqua 6→ x2 $H_2O$ 0.5→ 4% UA overnight 4°C, 2 h 50°→ SEM | Suppl.Fig.1i Fig.1e |
| 8 | Extending pyrogallol | Os 3→FeCN 17 → Os 3 → CaC 0.5→x2 $H_2O$ 0.5→ Pyrogallol 1.5/7/17→ x2 $H_2O$ 0.5→ Os-aqua 3/6→ µCT | Suppl.Fig.1j |
| 9 | Improving UA | Os 3→FeCN 17→ Os 3→ CaC 0.5→x2 $H_2O$ 0.5→ Pyrogallol 17→ x2 $H_2O$ 0.5→ Os-aqua 6→ x2 $H_2O$ 0.5→ 2% /4% UA overnight 4°C, 2 h 50°C→ µCT | Suppl.Fig.1 k Fig.1d |
| 10 | Cancelling Ld | Os 3→FeCN 17→ Os 3→ CaC 0.5→x2 $H_2O$ 0.5→ | Suppl.Fig.1 |

| | | | |
|---|---|---|---|
| | | Pyrogallol 17→ x2 $H_2O$ 0.5→ Os-aqua 6→ x2 $H_2O$ 0.5→ 4% UA overnight 4°C, 2 h 50°C→ Ld 4/24 50°C→µCT | k |
| 11 | Os very long | Os 3 days→ SEM<br>Os 6 days→ SEM | Suppl.Fig.2 c,d |
| 12 | Os 4°C vs. RT | Os 7 days 4°C→ SEM<br>Os 6 days 4°C, 1 day RT→SEM | Suppl.Fig.2 e,f |
| 13 | Os 4°C then FeCN | Os 6 days 4°C, 1day RT→FeCN 1 day 4°C/RT→SEM<br>Os 7 days 4°C→FeCN 1 day 4°C/RT→SEM | Suppl. Fig.2i,j |
| 14 | Os 4°C on hemisphere | Os 4°C → uCT at 17/24/40h | Suppl.Fig.3 a |
| 15 | CaC washing removes Os-FeCN redox gradient | Os 48→FeCN48→SEM<br>Os48→CaC48→FeCN48→SEM | Suppl.Fig.3 b,c |
| 16 | CaC washing speed up FeCN diffusion | Os 24 → FeCN 1.5 → uCT, SEM<br>Os 24 → CaC 24 → FeCN 1.5 → uCT, SEM | Suppl.Fig.3 d,e |
| 17 | Pg interaction with Os-FeCN redox gradient (H3) | Os 4°C 96, RT 24→CaC 4°C 48 →FeCN 4°C 48 (uCT) → Os 72→CaC 24(uCT) → $H_2O$ 29 (uCT) → Pg 24 (uCT) → $H_2O$ 48 → Os 48 → 4% UA 4°C 48, 50°C 5 →$H_2O$ 42 →SEM | Suppl.Fig3g |
| 18 | Pg interaction with Os-FeCN redox gradient and CaC (H1) | Os 4°C 63, RT 24 →CaC 4°C 24 → FeCN 4°C 48 → CaC 24 (uCT)→ Os 32→CaC 17 (uCT)→ $H_2O$ 9(uCT) → Pg 48 (uCT)→ $H_2O$ 40 → Os 48 → 4% UA 4°C 48, 50°C 5→$H_2O$ 48→ SEM | Suppl.Fig.3 g |
| 19 | CaC RT 2d removes Os-FeCN redox gradient (H4,5,6) | Os 4°C 72, RT 24→CaC 48 → FeCN 4°C 4 → CaC 24 (uCT H5) → Os 48→CaC 24→ $H_2O$ 24 (uCT H5) → Pg 24 (uCT H5) → $H_2O$ 24 → Os 24 (uCT H5) → 4% UA 4°C 48, 50°C 5 →$H_2O$ 48 →SEM | Suppl.Fig.3 h; Fig.2; Suppl.Fig.4; Suppl.Fig.7 |
| 20 | CaC washing at 4°C 4 day removes Os-FeCN redox gradient (H13) | Os 4°C 68 →CaC 4°C 96 → FeCN 4°C 72 → CaC 48 → Os 48 →CaC 24 → $H_2O$ 24 (uCT) → Pg 24 (uCT) → $H_2O$ 24 → Os 41 (uCT at 24/41 h) → 4% UA 4°C 48, 50°C 12 →$H_2O$ 24 →SEM | Suppl.Fig.3 h; Suppl.Fig.4 |

| | | | |
|---|---|---|---|
| 21 | Whole brain staining (W1,W2) | W1: Os 4°C 96 (uCT) → CaC 4°C 168→ FeCN 4°C 72 (uCT) → CaC 4°C 48, RT 48 → Os 48→CaC 72 (uCT) → $H_2O$ 48 (uCT) → Pg 24 (uCT) → $H_2O$ 48→ Os 96 (uCT) → 4% UA 4°C 48, 50°C 5.5 →$H_2O$ 24 →SEM<br><br>W2: Os 4°C 96 → CaC RT 72, 4°C 72→ FeCN 4°C 72 → CaC 4°C 48, RT 48 → Os 48→CaC 72 → $H_2O$ 48 (uCT) → Pg 24 → $H_2O$ 48→ Os 96 → 4% UA 4°C 48, 50°C 5.5 →$H_2O$ 24 →SEM | Fig.3; Suppl.Fig.7 g |
| 22 | 2nd Os stabilize samples in $H_2O$ | Os 24 → CaC 24 → FeCN 24 → Os 3 → CaC 0.5 → x2 $H_2O$ 0.5 → $H_2O$→ take LM images at different time points<br><br>Os 24 → CaC 24 → FeCn 24→ CaC 0.5 → x2 $H_2O$ 0.5 → $H_2O$ → take LM images at different time points | Supp.Fig.3i |
| 23 | Os vi staining | 1% Os (vi) CaC 6→ SEM | Suppl.Fig.5 c |
| 24 | Adding $CaCl_2$ in $OsO_4$ then FeCN | Os 24 (with $CaCl_2$)→ FeCN 17→ SEM | Suppl.Fig.5 g |
| 25 | $OsO_4$ at 4°C then FeCN | Os 24 4°C→FeCN17→SEM | Suppl.Fig.5 h |
| 26 | rO short vs. long | rO 3 → SEM<br>rO 24 → SEM | Suppl.Fig.5 e,f |
| 27 | CaC wash between $OsO_4$ and FeCN | Os 3/24→x3 CaC 1→FeCN 17→SEM | Suppl.Fig.5 a,b |

### Supplementary Figures

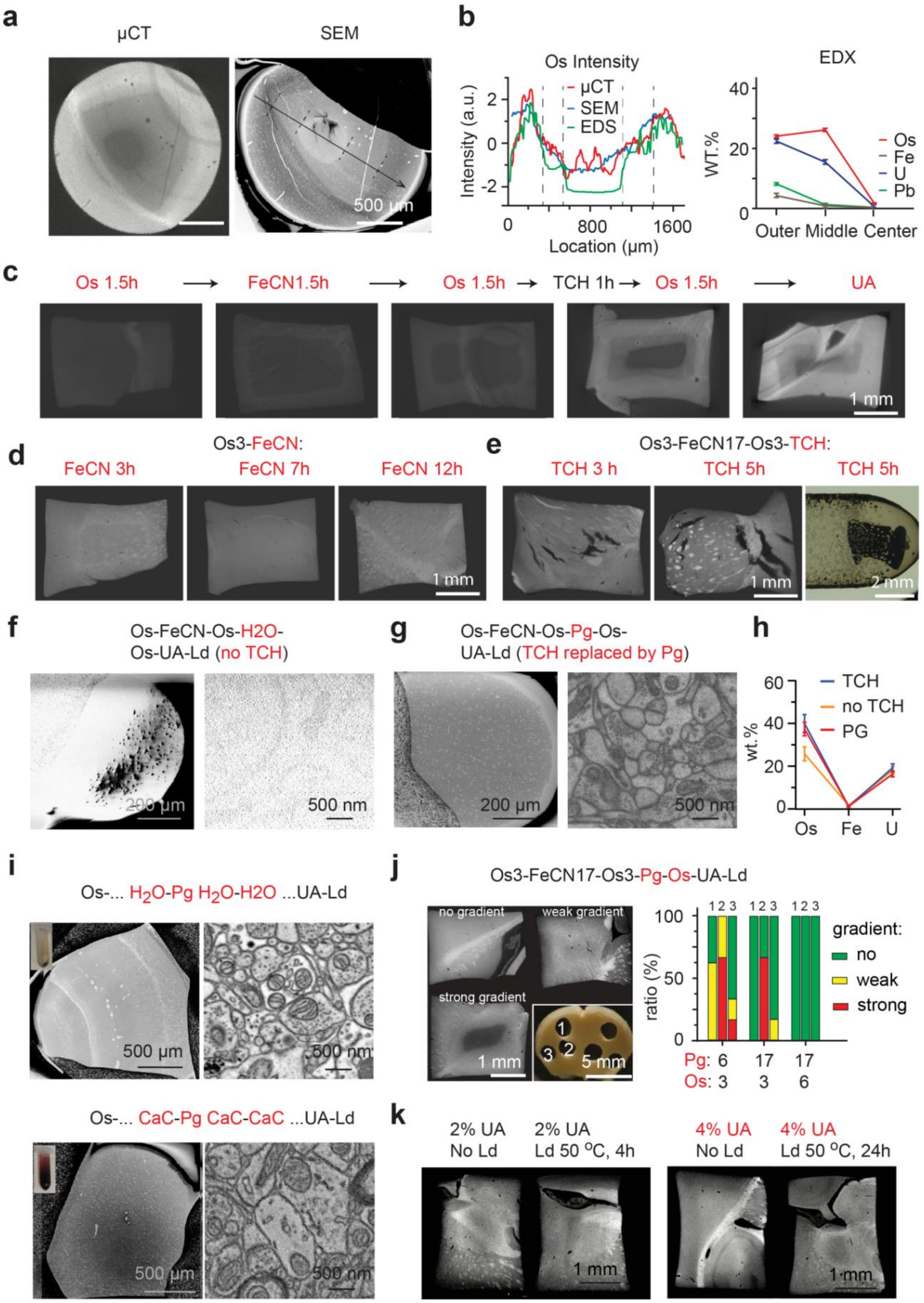


**Supplementary Figure 1. Analysis of staining gradients for 2-3 mm samples. (a)** Left: µCT image of staining gradient produced on 2 mm samples with Hua protocol(Hua et al., 2015) and confirmation with

SEM (right). **(b)** EDS analysis of staining gradient from sample in (a). **(c)** µCT diagnosis of staining gradient after each consecutive step of the Hua staining protocol(Hua et al., 2015) indicating that gradients occurred after FeCN and TCH steps. **(d)** Extension of FeCN step from 3 h to 12 h removed FeCN-related gradient. **(e).** Extending TCH incubation caused substantial breakages in the samples. **(f)** Omission of TCH step yielded very low sample conductivity; no meaningful images could be acquired in high vacuum SEM due to sample charging. **(g)** Replacing TCH by pyrogallol (PG (Mikula & Denk, 2015)restored sample conductivity. **(h)** EDS analysis showing that with PG incubation, the final osmium concentration in the samples reached the same level as using TCH. **(i).** Comparison of pyrogallol-related steps in water (top) vs. in CaC (bottom). When CaC buffer was used, the final contrast was decreased; a gradient was also observed. Electron dose for high resolution EM images in f,g,i:18 $e^-/nm^2$**.** **(j)** Effect of duration of pyrogallol related steps. Left: pyrogallol related gradient was dependent on location of tissue sample, with myelin-rich subcortical regions yielding stronger gradients; right: extension of pyrogallol incubation and 3$^{rd}$ $OsO_4$ step removed gradient for all sampling locations. **(k)** Gradients related to UA and Ld incubation. UA –related gradient could be removed by increasing concentration from 2% to 4%. Ld gradient could not be removed due to limited solubility and was omitted.

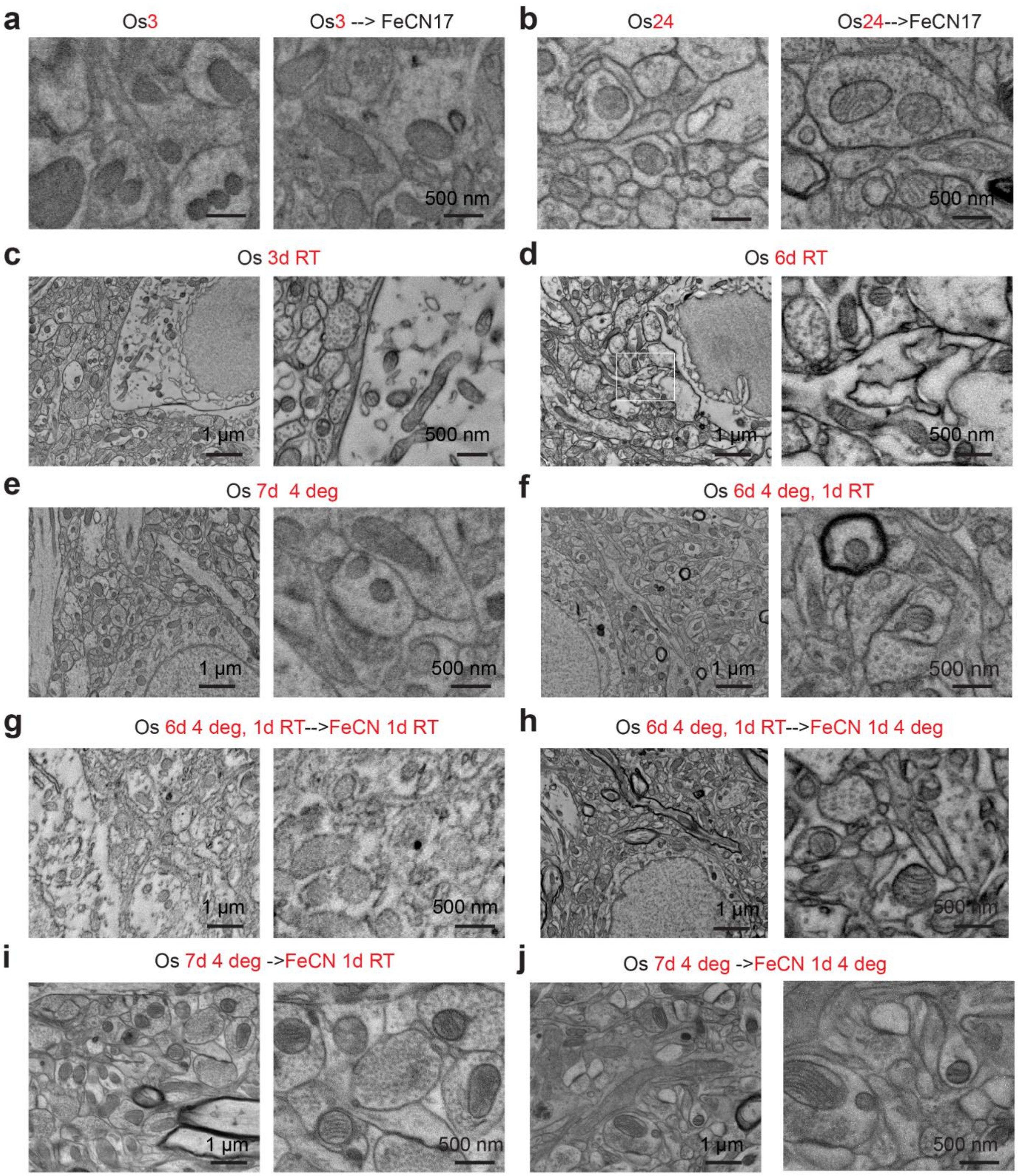


**Supplementary Figure 2. Effect of Os and FeCN incubation time and temperature on membrane contrast in 2-3 mm samples. (a)** Membrane contrast was not enhanced in samples stained with 3 h of $OsO_4$ (left) or with 3 h $OsO_4$ followed by 17 h of FeCN. **(b)** Membrane contrast was not enhanced in samples stained with 24 h of $OsO_4$ (left), but was enhanced in samples stained 24 h of $OsO_4$ followed by 17 h of FeCN. **(c,d)** 3 (c) and 6 (d) days of $OsO_4$ at room temperature yielded increased membrane contrast, respectively, but also substantial background extractions. **(e)** 7d incubation of $OsO_4$ at 4 °C avoided background extraction but yielded less contrast; **(f)** 6d incubation of $OsO_4$ at 4 °C followed by 1 d incubation at RT avoided background extraction and yielded better contrast. **(g).** 6d incubation of $OsO_4$ at

4 °C followed by 1 d incubation at RT, plus 1 day FeCN incubation at RT yielded destroyed ultrastructure; **(h)** 6d incubation of $OsO_4$ at 4 °C followed by 1 d incubation at RT, plus 1 day FeCN incubation at 4°C restored ultrastructural quality. **(i)** Membrane contrast enhancement also observed in samples stained for 7 days with $OsO_4$ at 4°C followed by 1 day of FeCN at RT. **(j)** Membrane contrast enhancement also observed in samples stained for 7 days with $OsO_4$ at 4°C followed by 1 day of FeCN at 4°C. All images were acquired in low vacuum SEM at 30 Pa, with maximum electron dose 70 $e^-/nm^2$.

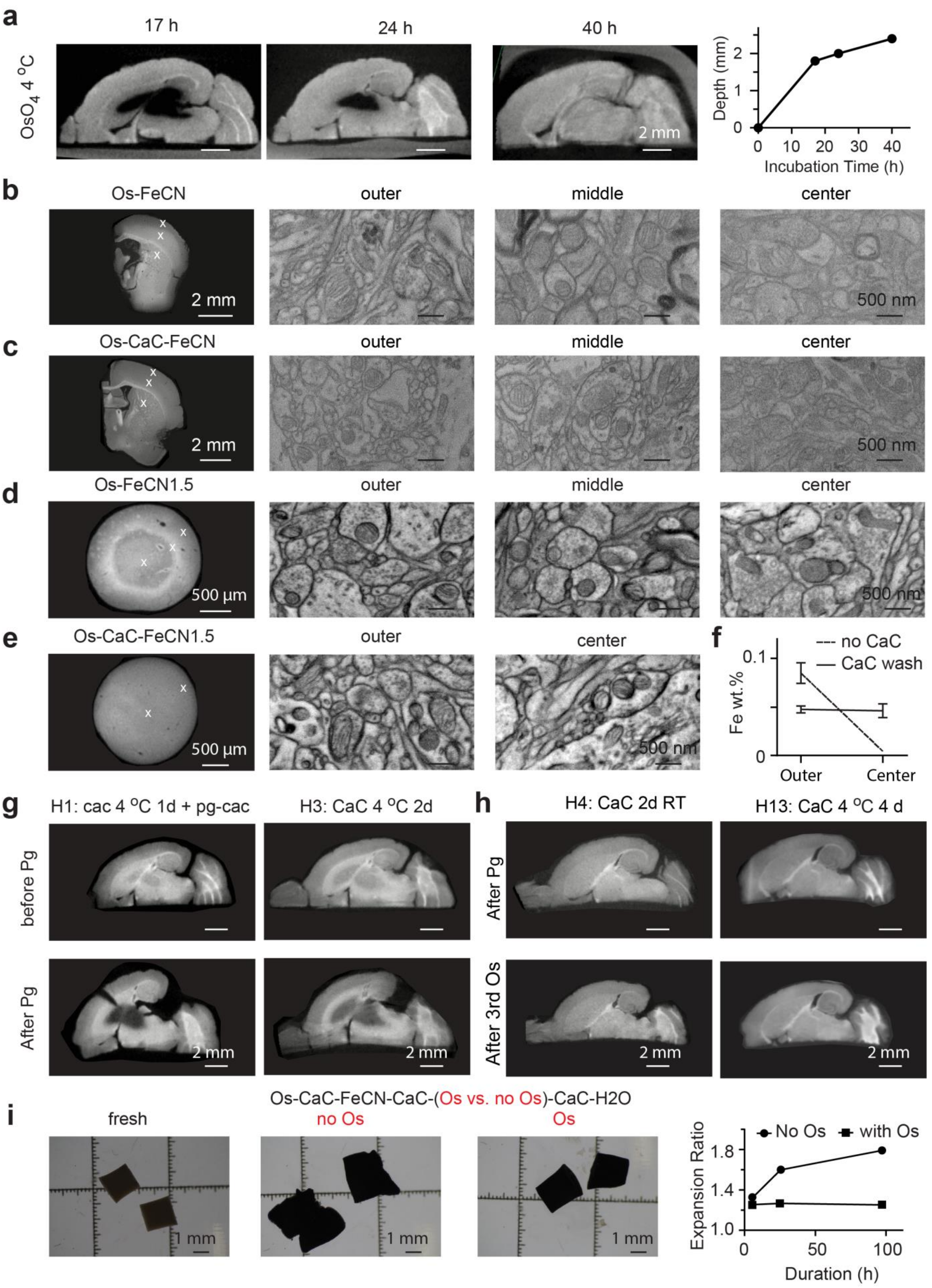

a
17 h
24 h
40 h
OsO4 4 °C
2 mm
Depth (mm)
Incubation Time (h)
b
Os-FeCN
outer
middle
center
2 mm
500 nm
c
Os-CaC-FeCN
outer
middle
center
2 mm
500 nm
d
Os-FeCN1.5
outer
middle
center
500 µm
500 nm
e
Os-CaC-FeCN1.5
outer
center
500 µm
500 nm
f
--- no CaC
— CaC wash
Fe wt.%
Outer
Center
g
H1: cac 4 °C 1d + pg-cac
H3: CaC 4 °C 2d
before Pg
After Pg
2 mm
h
H4: CaC 2d RT
H13: CaC 4 °C 4 d
After Pg
After 3rd Os
2 mm
i
Os-CaC-FeCN-CaC-(Os vs. no Os)-CaC-H2O
fresh
no Os
Os
1 mm
No Os
with Os
Expansion Ratio
Duration (h)

**Supplementary Figure 3. Gradients and breakages in mouse hemispheres. a** Observation of $OsO_4$ diffusion into mouse hemisphere sample at successive time points at 4°C using µCT. **b** $OsO_4$-FeCN incubation yielded macroscopic double-gradient with staining intensity decrease at the outer and innermost part of the sample. **c** Buffer rinsing step between $OsO_4$ and FeCN avoided the $OsO_4$-FeCN double gradient. **d** Except for generating double gradient, $OsO_4$-FeCN incubation slows down FeCN diffusion, causing no membrane contrast enhancing in the sample center. **e** Buffer rinsing step between $OsO_4$ and FeCN speed up the FeCN diffusion, lead to enhanced membrane contrast in sample center. **f** EDS measurement of the center of samples from d,e show the penetration depth of FeCN in c,d. **g** If the buffering step is 1 or 2 days at 4 °C, the OsO4 could not be easily washed out and would cause gradient (top), which would be amplified by Pg and also cause breakages (bottom) (note also in H1, CaC interaction due to not long enough $H_2O$ rinsing before Pg). **h** If the buffering step is 2 days RT or 4 days 4°C, the $OsO_4$ could be washed out and gradient would occur neither before Pg (top) nor after Pg (bottom). **i** Adding an extra $OsO_4$ incubation step after FeCN increases the sample stability in $H_2O$ for up to 100 h. All SEM images were acquired in low vacuum SEM at 30 Pa, with maximum electron dose 70 $e^-/nm^2$.

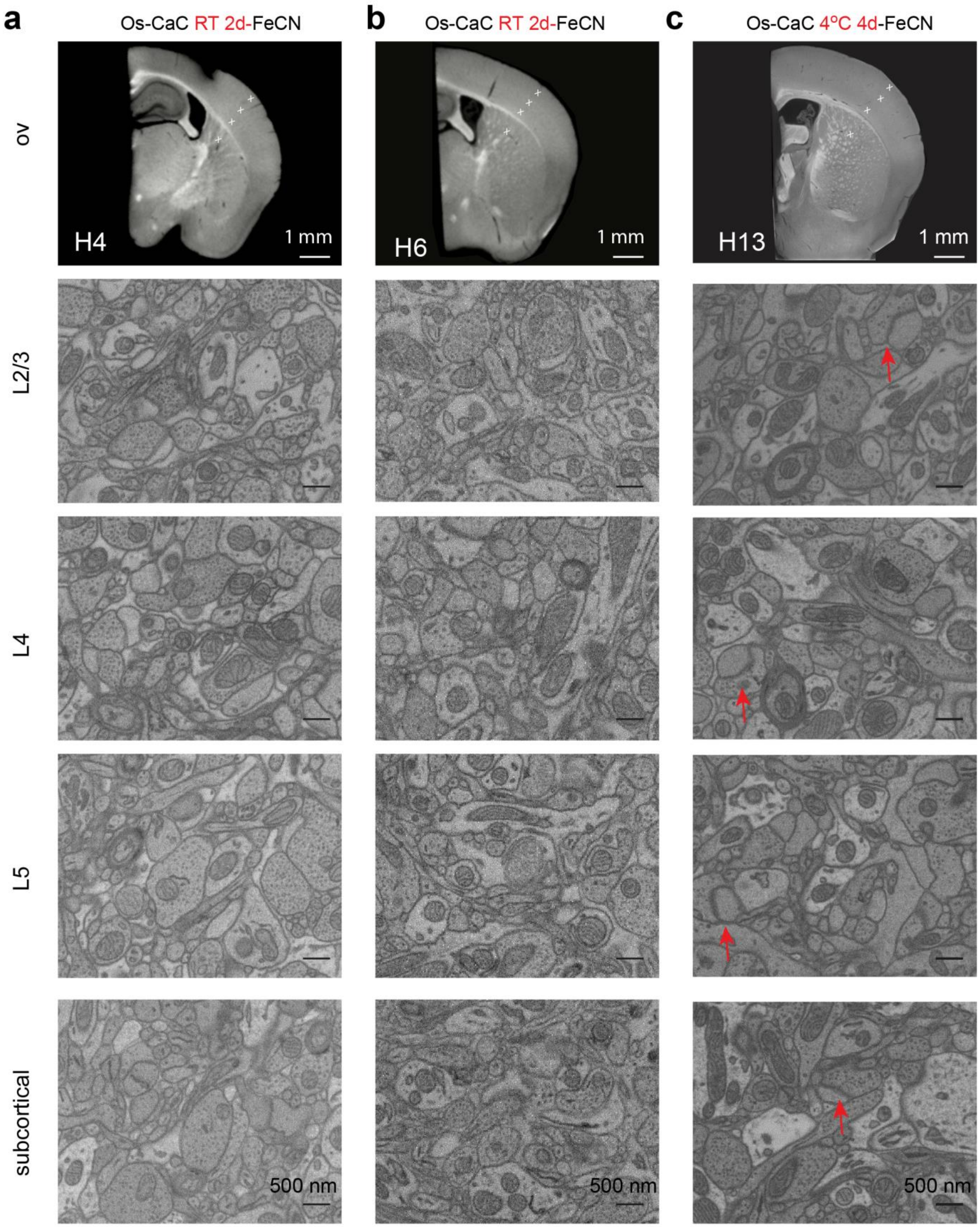


**Supplementary Figure 4. Reproduction of mouse hemisphere stainings. (a,b)** Hemispheres H4 and H6 stained as in Table 1, Suppl. Table 3. **(c)** Hemisphere H13 stained as in Table 1, Suppl. Table3. Together with H5 (Fig.2) and H13, a total of n=4 hemispheres were successfully stained with parameters reported in Table 1, Suppl.Table 3. Note also in H13, the PSD preservation is better. All SEM images were taken at high vacuum with a maximum dose of 84 $e^{-}/nm^{2}$.

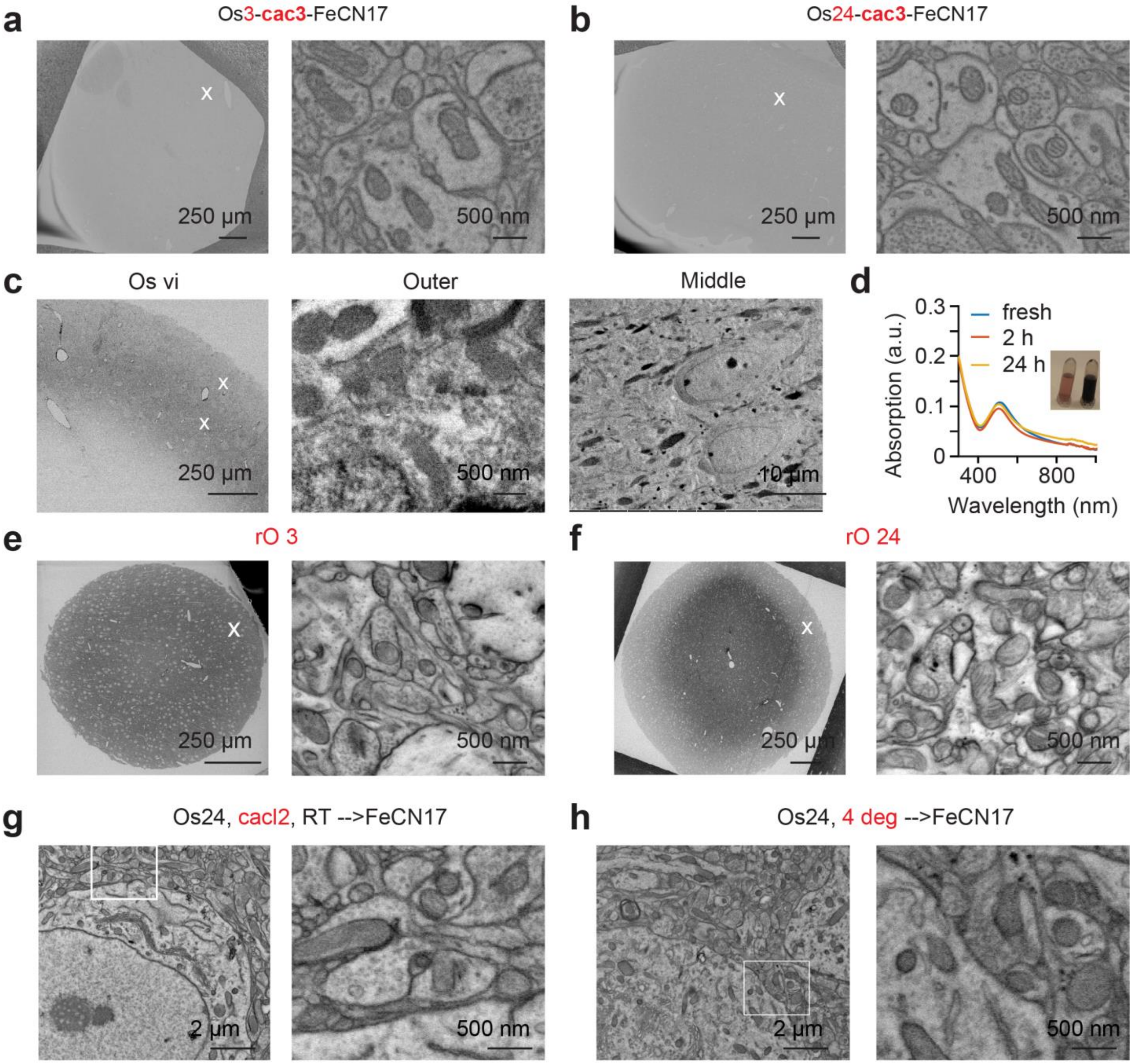


**Supplementary Figure 5. Experiments conducted for exploring possible chemical mechanisms underlying staining contrast generation. a,b** When adding an intermediate CaC washing step between $OsO_4$ and FeCN, the final contrast remained similar to 2 mm protocols (Fig.1c,d). Imaged in SEM in high vacuum, electron dose 18 $e^-/nm^2$. **c** Staining with potassium osmate (vi) yielded a precipitation band, however did not yield enhanced contrast. **d** UV-vis spectrum of potassium osmate (vi) in sodium cacodylate buffer over 24 h. **e,f** 2 mm samples stained with “reduced osmium” protocol for 3h vs. 24h. **g** adding $CaCl_2$ into $OsO_4$ abolished the membrane contrast enhancement of FeCN. **h** $OsO_4$ incubation at 4°C also abolished the membrane contrast enhancement effect of FeCN. SEM acquisition for panels c,e,f,g,h was performed in low vacuum at 30 Pa, maximum electron dose of 70 $e^-/nm^2$

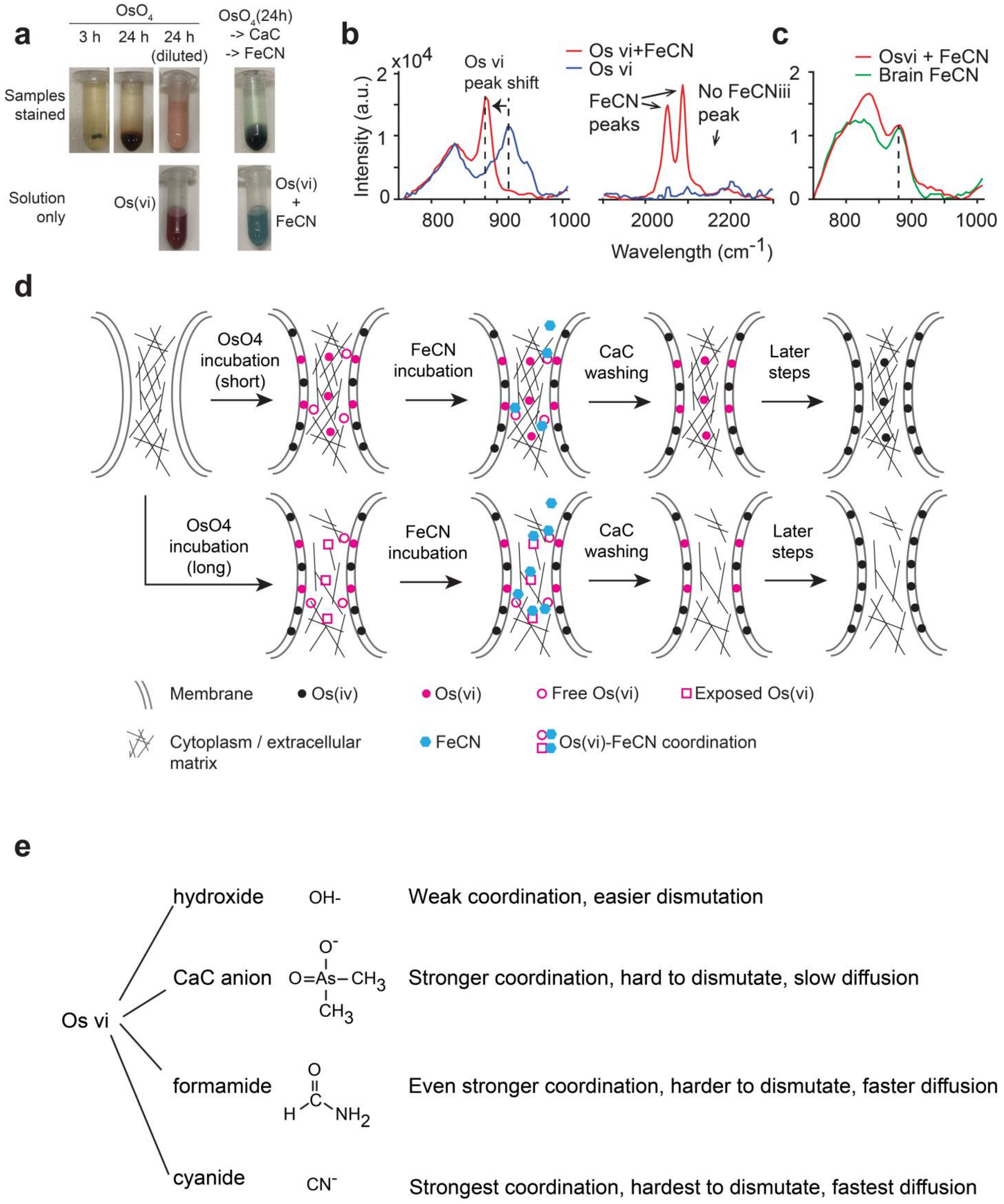


**Supplementary Figure 6. Possible chemical logic underlying staining results (see Suppl. Text and Suppl. Fig. 5). (a)** Comparison of bright-field color appearance of samples stained in $OsO_4$ for 2h, 24h, 24h (diluted to evaluate color), and stained in $OsO_4$(24h)->CaC->FeCN (intermediate CaC wash to remove $OsO_4$), top row. For comparison, pure staining solutions are shown below: Os(vi) and in vitro reaction of Os(vi) with FeCN (bottom). Note that sample solution after 24h OsO4 incubation appears similarly colored as Os(vi) solution (pink); and sample solution after Os-Cac-FeCN staining appears similarly colored as Os(vi)+FeCN reaction in vitro (blue-green). **(b). Left**: Raman spectrum measurements of the product of Os(vi) + FeCN showing a shifted Os(vi) peak, indicating potential coordination reaction between Os(vi) and FeCN; **middle**: in the wavelength range corresponding to FeCN(ii) and FeCN(iii), however, we only found signal consistent with FeCN(ii) peaks, but not FeCN(iii), suggesting the reaction between Os(vi) and FeCN was not a redox reaction. **(c)**. The same Os(vi) signal was observed in brain staining solution after FeCN staining and in-vitro Os(vi) + FeCN reaction. **(d)** Sketch summary of the possible chemical reactions. Short $OsO_4$ incubation may deposit Osmium into both membrane and background (primarily proteins in the cytosol). Os(vi) would be produced in the membrane and also within the extracellular matrix. While membrane-related Os(vi) would have access to the aqueous phase, protein-bound may be shielded from converting into free Os(vi). If FeCN was then applied to the sample, it would donate $CN^-$ as coordination ligand to Os(vi); however, this would only occur for free Os(vi) since Os(vi) in the background or membranes would not yet be as accessible for FeCN. This would keep osmium in both membranes and backgrounds, resulting in no contrast enhancement or slightly reduced contrast. But if the first $OsO_4$ incubation was prolonged, $OsO_4$ over-oxidization of proteins can cause the 3D conformational change of background proteins, and expose hidden Os(vi) to the aqueous phase. If FeCN was applied afterwards, FeCN would donate $CN^-$ to free Os(vi) and Os(vi) exposed within the cytosolic/ECM protein network. This would result in removal of background osmium and could thus increases local contrast. **(e)**  Summary of Os(vi) coordination chemistry potentially relevant for staining mechanisms.

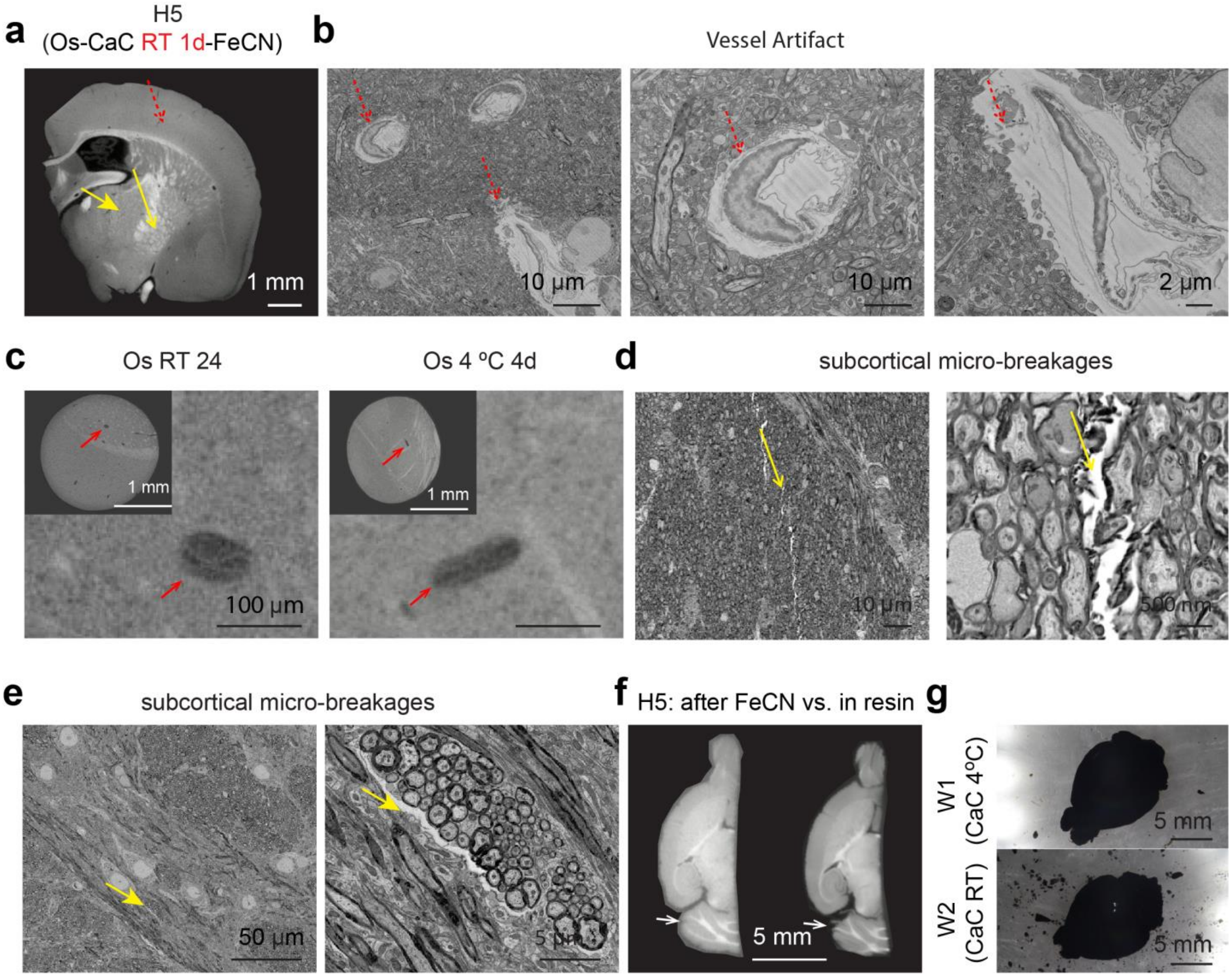


**Supplementary Figure 7.** Summary of remaining artifacts in whole-brain staining protocol. **(a)** µCT cross section of H5 showing locations with artifacts detailed in b-e. **(b)** Red arrow: "Vessel artifact": disruption of blood vessel pericytes from surrounding neuropil. **(c)** Vessel artifact visible in µCT images of 2 mm samples stained with $OsO_4$ RT 1 day or 4°C for 4 days. **(d,e)** Yellow arrows: Micro-breakages with width of less than 1 µm were occasionally observed in subcortical areas, in particular in highly myelinated regions. **(f)** During $H_2O$ incubation, the cerebellum was especially sensitive to macroscopic damage. **(g)** Changing the CaC rinsing step between $1^{st}$ $OsO_4$ and FeCN to 4°C improved the stability of the cerebellum for whole brain staining. All SEM images were acquired at high vacuum, with maximum dose of 21 $e^{-}/nm^2$.
